# Fos regulates age-dependent neuroinflammation in *VAPB^ALS^*

**DOI:** 10.64898/2026.02.28.708780

**Authors:** Namrata Pramod Kulkarni, Aparna Thulasidharan, Amarendranath Soory, Pulkit Goel, Vidyadheesh Kelkar, Girish S Ratnaparkhi

## Abstract

Amyotrophic Lateral Sclerosis (ALS) is a fatal neurodegenerative disorder characterized by progressive loss of motor function. We have developed a *Drosophila* model of ALS8 (*VAPB^P58S^*) using CRISPR/Cas9 genome editing. VAPB is an ER-based adapter protein associated with and regulating intracellular membrane:membrane contact sites.

*VAPB^P58S^* flies show progressive age-dependent motor deficits and a shortened lifespan, paralleling features of the human disease. *VAPB^P58S^* brains exhibit age-dependent neuroinflammation, as measured by whole-transcriptome quantitative mRNA sequencing, suggesting a broad, low-grade enhancement in signalling in multiple (Toll, IMD, Jak-STAT and Jun-kinase) immune pathways. We implicate glial cells in the brain as the site of brain inflammation and identify *Drosophila* Fos (Kayak) as a key modulator of age-dependent inflammation. In accordance, we find that overexpression of wild-type *kayak* or its dominant-active variant *kayak^K357R^* in glia reduces inflammation and, concomitantly, improves motor function. In contrast, knockdown of glial *kayak* accelerates age-dependent deterioration of motor function and enhances neuroinflammation.

Our study underscores the roles of glial-modulated brain inflammation in dictating ALS8 progression and identifies *kayak* as a central negative regulator of neuroinflammation in disease.

**Summary Statement:** We uncover definitive evidence for age-dependent neuroinflammation, originating from glial cells and regulated by Fos, as a key mechanism underlying Amyotrophic Lateral Sclerosis 8.

## Introduction

Amyotrophic Lateral Sclerosis (ALS) is a disease characterized by muscle atrophy owing to the degeneration of neurons in the cortex, brainstem and spinal cord (Feldman et al., 2022; Mitchell and Borasio, 2007; Pasinelli and Brown, 2006). The disease starts with mild symptoms like weakness, difficulty swallowing and slurred speech, but quickly progresses to more severe symptoms like motor dysfunction, labored breathing and paralysis. Death typically occurs due to respiratory failure within 3-5 years of diagnosis (Boylan, 2015; Cleveland and Rothstein, 2001; Tartaglia et al., 2007). 90% of the ALS cases are sporadic ALS (sALS) with no identified family history, whereas the remaining 10% are classified as familial ALS (fALS) involving one or more affected family members (Boylan, 2015; Marin et al., 2012). More than 30 loci contributing to distinct or overlapping pathomechanisms, such as oxidative stress, neuroinflammation, lipid dysregulation, mitochondrial dysfunction, and proteostasis failure, have been associated with ALS, indicating the multifactorial and complex nature of the disease (Abel et al., 2012; Jiang et al., 2024; Robberecht, 2000; Tendulkar et al., 2022; Thulasidharan et al., 2024).

Although ALS was first described almost a century ago, little has been achieved in terms of treatment, with very few available drugs, offering only modest benefits (Paganoni et al., 2020; Wijesekera and Leigh, 2009; Yoshino and Kimura, 2006). Additionally, due to its oligogenic nature, the molecular mechanism underlying ALS remain poorly understood, rendering the disease largely untreatable. Thus, efforts to investigate molecular mechanisms and gene networks underlying ALS in model organisms can pave the way for the development of more effective therapies.

The eighth ALS locus to be identified was vesicle-associated membrane protein (VAMP) associated protein B (*VAPB*). A dominant proline-to-serine point mutation at position 56 (P56S) in the MSP domain was identified in Portuguese-Brazilian families and is associated with fALS (Nishimura et al., 2004). *VAPB* is a definitive ALS locus (Abel et al., 2012; McCann et al., 2020). It encodes a type II integral membrane protein that resides on the endoplasmic reticulum (ER) membrane. VAPB has three domains: the C-terminal transmembrane domain (TMD), which is inserted into the ER membrane; a coiled-coil domain (CCD) that helps form multimers; and the N-terminal major sperm protein (MSP) domain, which faces the cytosol (Borgese et al., 2021; James and Kehlenbach, 2021; Kors et al., 2022). The MSP domain can interact with a wide range of partner proteins that decorate the surface of organelle membranes and facilitate the formation of the membrane–membrane contact site (MCS) between the ER and these (other) organelles. VAPB, through the MCS, plays a crucial role in efficiently coordinating inter-organelle processes, thereby maintaining cellular homeostasis. These cellular processes include membrane trafficking, lipid transport & lipid homeostasis, immune signalling, calcium signalling, autophagy, stress response & protein homeostasis (reviewed in (Borgese et al., 2021; James and Kehlenbach, 2021; Murphy and Levine, 2016; Neefjes and Cabukusta, 2021; Reinisch et al., 2025; Zhao et al., 2018)). Modulation and disruption of VAP-related MCS can affect these functions, in whole or in part (Kodama et al., 2025; Obara et al., 2024; Stoica et al., 2014).

Our laboratory has been interested in studying the *Drosophila* orthologue of *VAPB*, *Vap33/CG5014* (hereafter referred to as *VAP*), as a model for ALS8. As a first step, we used an enhancer-suppressor screen to identify the VAP gene regulatory network (GRN) in flies. The study uncovered multiple ALS-ortholog loci, such as *superoxide dismutase 1* (*SOD1*), *Alsin*, *TAR DNA-binding protein 43* (*TDP43*), and *Target of Rapamycin* (TOR), as genetic interactors of *VAP* (Deivasigamani et al., 2014). Subsequently, we studied the effects of SOD1 and TOR modulation on VAP^P58S^ aggregates; *VAP^P58S^* is the *Drosophila* equivalent of *VAPB^P56S^*. Notably, our study revealed that in the larval brain, reducing either SOD1 levels or TOR signalling increased cellular ROS levels, which, in turn, enhanced clearance of VAP^P58S^ aggregates by triggering the ubiquitin-proteasome system (UPS). Further, using a null-rescue model developed by the Tsuda lab, *gVAP^P58S^* (Moustaqim-Barrette et al., 2014), we discovered that autophagy, rather than UPS was the dominant mechanism contributing to aggregate clearance (Thulasidharan et al., 2024) in the adult brain. Additionally, we found that age-dependent inflammation was a key factor in determining the extent of disease progression, with Immune Deficient (IMD) signalling upregulated in *gVAP^P58S^* (Tendulkar et al., 2022).

In the current study, we aimed to extend our understanding of the roles of neuroinflammation in disease pathogenesis, using a CRISPR-Cas9 genome-edited *VAP^P58S^* model generated in our laboratory. As a first step, we conducted an age-dependent quantitative transcriptomic experiment on the heads of the CRISPR-generated *VAP^P58S^* mutant line. Interestingly, the *VAP^P58S^* transcriptome showed a low-grade increase in the levels of inflammatory markers with age. Intriguingly, our analysis revealed that the neuroinflammatory changes were global and not restricted to a single pathway, with inflammatory markers detected from the IMD, Toll, c-Jun N-terminal kinase (JNK), and Janus Kinase-Signal Transducer and Activator of Transcription (Jak-STAT) pathways. We found that in *VAP^P58S^*flies, glial expression of the *Drosophila* ortholog of Fos, *kayak* (*kay*), reduced transcription of inflammatory markers and, concomitantly, slowed the progression of age-dependent motor defects. Our study highlights the role of Kay as a broad, critical modulator of neuroinflammation and disease progression and functionally links VAP, located in membrane:membrane contact sites to signalling by innate immune pathways.

## Results

### A genome-edited *VAP^P58S^*model for ALS8

The initial fly disease models of *VAP^P58S^* relied on overexpression (OE) of the transgene (Han et al., 2012; Mitne-Neto et al., 2011; Pennetta et al., 2002; Ratnaparkhi et al., 2008; Tsuda et al., 2008) and the subsequent assessment of neuronal physiology. Two weaknesses of the OE model were that the mutant VAP^P58S^ was present in excess, and endogenous VAP^WT^ was present. Subsequently, Hiroshi Tsuda’s laboratory developed an improved model in which an endogenous *genomic-VAP* (*gVAP^WT^* or *gVAP^P58S^*) insert on the third chromosome was used to rescue the lethality of the VAP null (*ΔVAP*) (Moustaqim-Barrette et al., 2014). The Tsuda disease model, *ΔVAP;gVAP^P58S^*, unlike its control *ΔVAP;gVAP^WT^*, has a shorter lifespan and shows progressive motor dysfunction. The *ΔVAP;gVAP^P58S^* model has VAP^P58S^ inclusions in the brain; with cells showing ER expansion and ER stress (Moustaqim-Barrette et al., 2014; Tendulkar et al., 2022; Thulasidharan et al., 2024). The Tsuda model did not exhibit the artefacts observed in the OE models, though it was constrained to studying only males. Additionally, two chromosomes (I & III) were occupied, limiting complex genetic manipulations.

Here, we have generated a *VAP^P58S^* mutant line by editing the wild-type *VAP* locus using CRISPR-Cas9 genome editing (Suppl. Fig. 1). Details of the editing experiment are provided in the Materials and Methods section, with a summary included in Suppl. Fig. 1. The *VAP^P58S^* genome-edited line (*VAP^P58S^* hereafter) is homozygous viable with a lifespan comparable to *ΔVAP;gVAP^P58S^*. We find that both *VAP^P58S^* mutant males (Fig. 1A, red curve) and females (Fig. 1C, red curve) showed a significant reduction in survival as compared to the controls (*VAP^WT^*; Fig. 1A and 1C, black curve, respectively). *VAP^P58S^* males have a median lifespan (ML) of 22 days, while females show a ML of 21.5 days, which is significantly less than the MLs of 44 and 50 days for *VAP^WT^* males and females (Fig. 1B and 1D), respectively. Compared with *VAP^P58S^*, the *ΔVAP;gVAP^P58S^* males have an ML of 27 days. The *VAP^WT^* animals used were ‘CRISPR’ controls; flies that had been part of the genome-editing pipeline but did not acquire the P58S mutation in the *VAP* locus.

**Figure 1:**
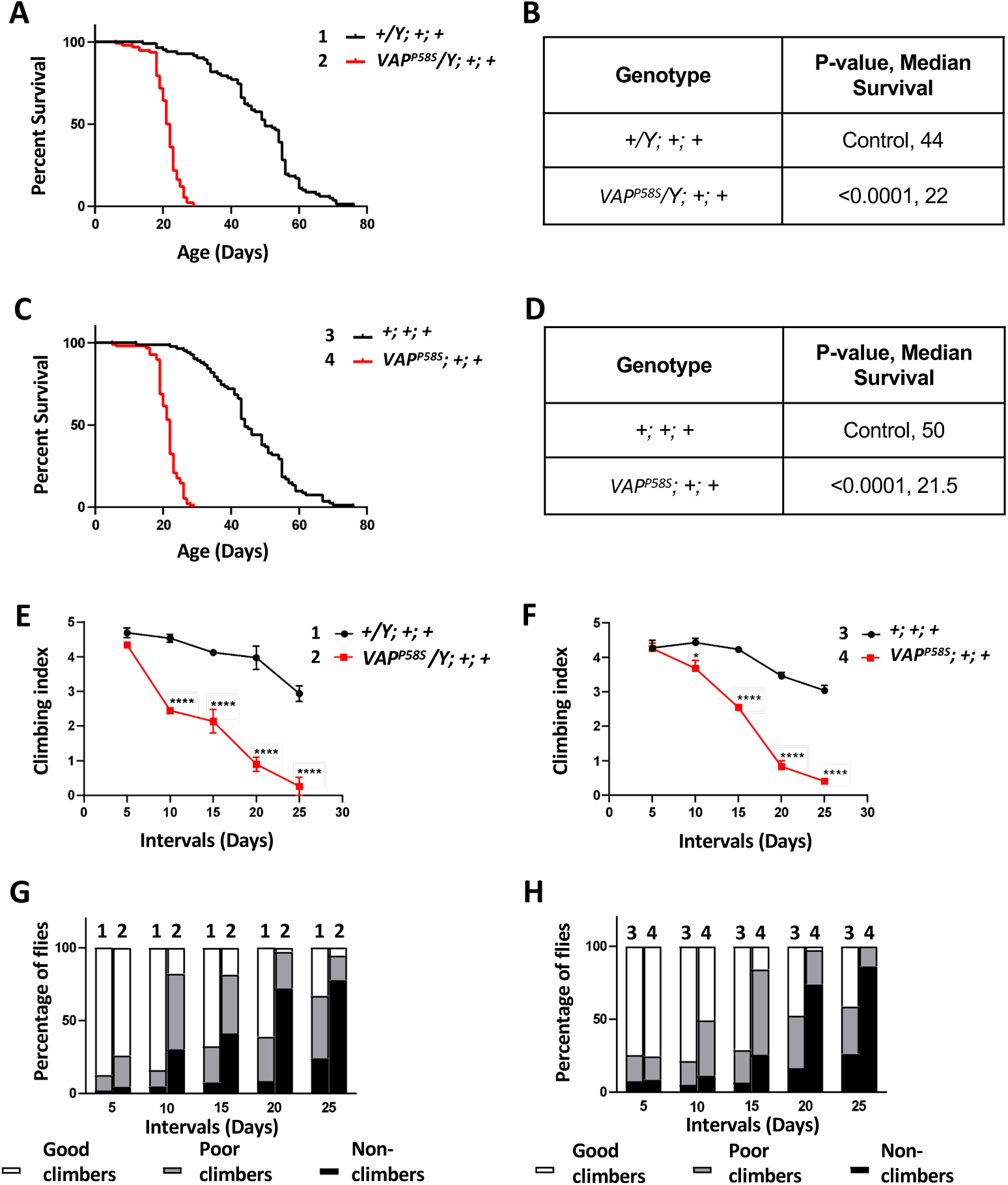
The *VAP^P58S^*mutant has reduced survival and exhibits progressive age-dependent motor defects as compared to *VAP^WT^*. **A.** Lifespan curves for *VAP^P58S^* males (2, red curve) & *VAP^WT^* (1, black curve). n=80-100 flies for every genotype, N= 3. Curve comparison was done using Log-rank test. Combined p-value for the set is <0.0001. **B.** Tabulated results for the lifespan in panel A, quantified in terms of ML. **C.** Lifespan curve for *VAP^P58S^* females (4, red curve) & *VAP^WT^* (3, black curve). n=80-100 flies for every genotype, N=3. Curve comparison was done using Log-rank test. Combined p-value for the set is <0.0001. **D.** Tabulated results for the lifespan in panel C, quantified in terms of ML. **E.** Climbing index for *VAP^P58S^* males (2, red curve) plotted for intervals 5, 10, 15, 20 and 25 days. *VAP^WT^* (1, black curve) was used as the control. n=25-30, N=3 flies for every genotype. Day 5, p> 0.05 (ns), Day 10, p<0.0001 (****), Day 15, p<0.0001 (****), Day 20, p<0.0001 (****), Day 25, p<0.0001 (****). **F.** Climbing index for *VAP^P58S^*females (4, red curve) plotted for intervals 5, 10, 15, 20 and 25. *VAP^WT^*(3, black curve) was used as the control. n=25-30, N=3 flies for every genotype. Day 5, p> 0.05 (ns), Day 10, p<0.05 (*), Day 15, p<0.0001 (****), Day 20, p<0.0001 (****), Day 25, p<0.0001 (****). **G.** Percentage of good climbers (white), poor climbers (grey) and bad climbers (black) for *VAP^P58S^* males. *VAP^WT^* was used as the control. **H.** Percentage of good climbers (white), poor climbers (grey) and bad climbers (black) for *VAP^P58S^* females. *VAP^WT^* was used as the control.

### *VAP^P58S^* mutant flies show a progressive increase in age-dependent motor defects

To characterise motor defects in the *VAP^P58S^* line, we performed an age-dependent startle-induced negative geotaxis assay (Madabattula et al., 2015; Tendulkar et al., 2022). The *VAP^P58S^* mutant flies, both males and females showed no obvious motor defects at eclosion. With age, both male and female *VAP^P58S^*flies showed deterioration in climbing ability. The climbing assay was performed on days 5, 10, 15, 20 and 25 for both males and females. *VAP^P58S^* mutant males showed no motor defects on day 5 when compared to the *VAP^WT^,* but showed a significant decline in the climbing index by day 10, which further worsened with age. The *VAP^WT^*males, on the other hand, showed a mild decline in motor abilities as age progressed (Fig. 1E).

Similarly, *VAP^P58S^* mutant females showed no motor defects on day 5 when compared to the age-matched *VAP^WT^* females, but showed a progressive deterioration in motor function with age. *VAP^WT^* females, on the other hand, showed a mild decline in motor abilities with age (Fig. 1F). Both male and female *VAP^P58S^* mutant lines became completely immobile by day 25. Also, both male and female *VAP^P58S^* mutant lines showed a decrease in the number of good climbers and a corresponding increase in the number of poor climbers and non-climbers as they aged. Eventually, the poor climbers converted to non-climbers (Fig.1G and 1H, respectively). Thus, we conclude that *VAP^P58S^* mutant flies exhibit progressive, age-dependent motor defects, and their behavioural phenotypes are comparable to those observed in the existing *ΔVAP;gVAP^P58S^* disease model (Moustaqim-Barrette et al., 2014).

### *VAP^P58S^* flies show enhanced age-dependent neuroinflammation

In a previous study (Tendulkar et al., 2022) using the *ΔVAP;gVAP^P58S^* model of *ALS8*, we uncovered a role for the IMD– nuclear factor kappa beta (NFκB) pathway in disease progression, with Caspar OE in glia reducing neuro-inflammation and improving motor function. In this study, to thoroughly quantify the onset of inflammation in the adult brain, we measured the head transcriptome in ageing *VAP^P58S^* mutant flies, with *VAP^WT^*animals used as age-matched controls. Transcriptomes were measured at days 5, 15 and 20 in the heads of *VAP^P58S^* and *VAP^WT^* males, based on lifespan and motor-function data (Fig. 2). The *VAP^P58S^* line enabled us to collect data from both males and females (Suppl. Fig. 3), allowing us to address sexual dimorphism in our disease model.

**Figure 2:**
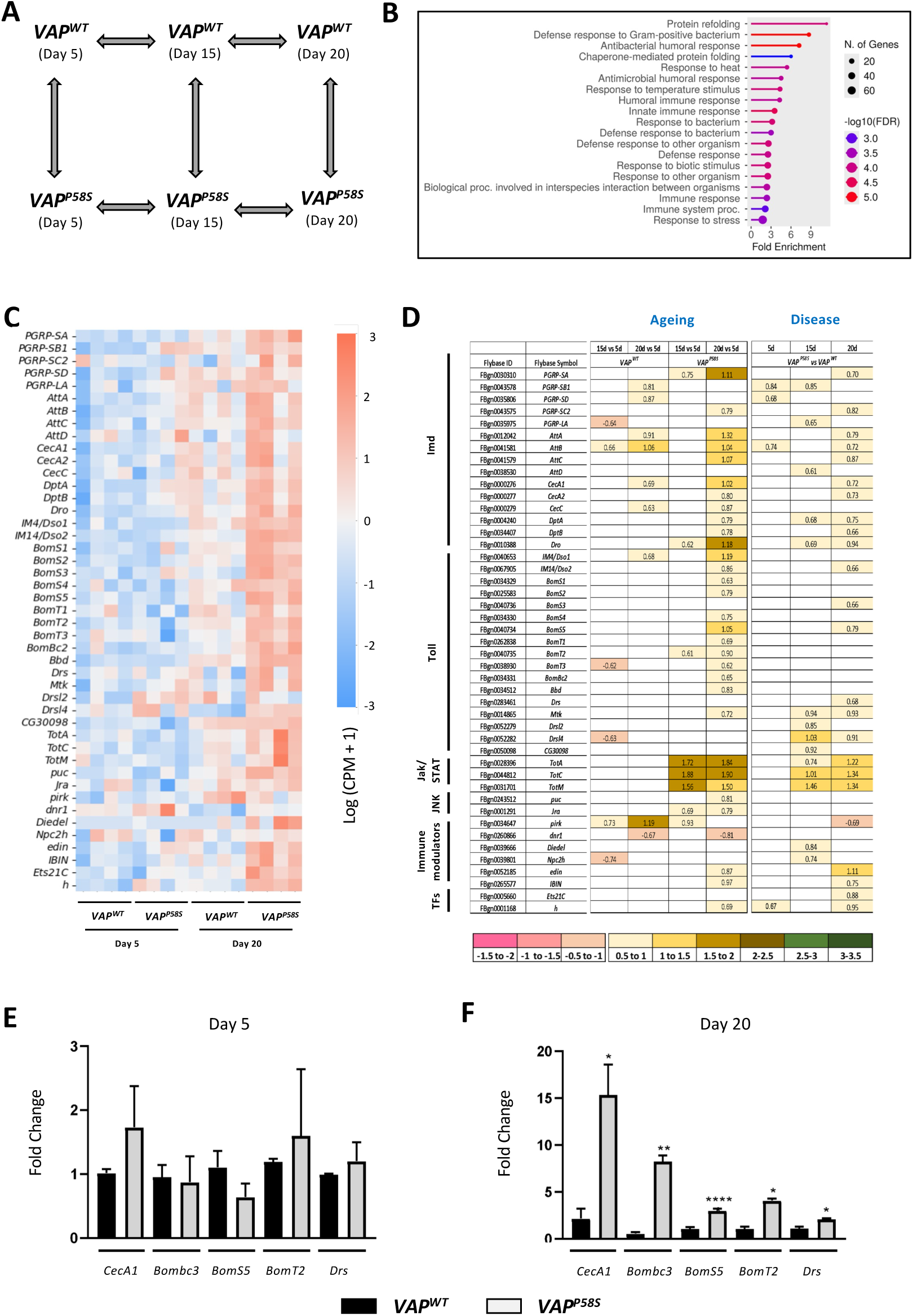
Transcriptomic study on *VAP^P58S^* male heads showing increased neuroinflammation with age. **A.** Genotypes and timepoints used for the 3’ mRNA sequencing and the pairwise comparison done to obtain Differentially Expressed Genes (DEGs). **B.** Gene Ontology (GO) enrichment analysis for significantly differentially expressed genes between *VAP^P58S^* and *VAP^WT^* at days 5, 15 and 20 combined. **C.** Heatmap showing the normalized expression counts for immune genes significantly differentially expressed between *VAP^P58S^* and *VAP^WT^* for days 5 and 20. **D.** Tabular representation of log_2_FC values for immune gene transcripts. The genes have been categorized into their immune pathway or the molecular function they perform. ‘Ageing’ depicts a comparison of the genotype with itself as it ages, while ‘Disease’ shows a comparison of *VAP^P58S^*with *VAP^WT^* at age-matched timepoints. **E. & F.** qRT-PCR for the Imd pathway target *CecA1* and Toll pathway targets *Bombc3*, *BomS5*, *BomT2* and *Drs* at days 5 and 20, respectively, for *VAP^WT^* and *VAP^P58S^* heads. The Y-axis shows log_2_FC normalised to *rp49*, which is a housekeeping gene. Values shown as mean + SEM. n (number of heads) = 40-50, N (Biological replicate) = 3. Statistical analysis was done using two-way ANOVA. Multiple comparisons were performed using Tukey’s test. *p< 0.05, **p< 0.01, ***p< 0.001.

For females, data were collected for two time points, day 5 and day 20. Our experiments utilised the Lexogen QuantSeq kit to generate multiplexed libraries for measuring quantitative gene expression (described by (Soory and Ratnaparkhi, 2022) and also in Materials and Methods). A schematic of the experimental timepoints, genotypes, and pairwise comparisons is shown in Fig. 2A & Suppl. Fig. 3A, with Suppl. Fig 2A showing the 3’ mRNA sequencing workflow. Volcano plots for male and female data, with a few important genes highlighted, have also been plotted (Suppl. Fig 2B and 2C, respectively). Fig. 2 displays data for males. Gene Ontology (GO) analysis (Fig. 2B) of the differentially expressed genes between *VAP^P58S^* and *VAP^WT^* males indicates that the most significant changes in the transcriptome are associated with host defence response, with genes classified as ‘immunity’ markers upregulated. The immune response category is followed by a ‘Chaperones/Protein refolding’ category that includes Heat Shock proteins of the Hsp70, Hsp60 family, as well as small heat shock proteins, which are biomarkers of ageing (Tower, 2011). Both categories are expected, as research from our laboratory has highlighted both protein aggregation (Chaplot et al., 2019; Thulasidharan et al., 2024) and inflammation (Tendulkar et al., 2022) as key features of the *ΔVAP;gVAP^P58S^* model (Moustaqim-Barrette et al., 2014). What is surprising, however, is the dominance and preponderance of these genes in the transcriptomic landscape of the disease model. Of the ∼500 genes statistically upregulated in the *VAP^P58S^* model, ∼10% are immune genes, and ∼2.5% are protein-folding-related. Genes linked to neuronal function are also modulated, as discussed in a subsequent section.

To understand how the transcriptome changes across genotypes and with age, we plotted a heatmap for a subset of these differentially expressed genes (Fig. 2C). The immune genes, especially ‘defence genes’ showed increased expression in *VAP^P58S^*as compared to *VAP^WT^*, as well as age-dependent enhancement (15-day data excluded for clarity and is shown as Suppl. Fig. 2D). The heat map depicts normalized expression counts for immune genes, indicating weak to moderate changes in transcript levels. Nevertheless, the 2-6-fold increase in transcripts of defence genes at day 20 is striking, especially given that the *VAP^P58S^* at day 5 shows non-significant changes compared to age-matched *VAP^WT^*.

To further gain insights into the immune pathways contributing to this low-grade inflammation, we segregated defence genes into known immune signal transduction pathways and tabulated log_2_FC values for each transcript (Fig. 2D). A large number of defence genes represented are anti-microbial peptide (AMP) genes that are strong indicators of upregulation of specific immune-responsive signalling pathways. We categorised the genes that changed as ‘Ageing’ vs ‘Disease’. For ageing, we highlighted genes that showed a significant change in expression levels at 20- or 15-days vs 5 days, for both *VAP^P58S^* and *VAP^WT^*, independently. For disease, we compared age-matched *VAP^P58S^*vs *VAP^WT^* heads (5-day, 15-day or 20-day).

In the ‘Ageing’ context, both *VAP^WT^* and *VAP^P58S^* showed an increase in the expression of immune genes, however, *VAP^P58S^*showed higher log_2_FC values as compared to *VAP^WT^*, for a subset of genes (Fig. 2D). Transcripts with higher values included *Attacins (Att)*, *Cecropins (Cec)*, *Diptericins (Dpt)*, *Drosocin (Dro)*, *Daisho (Dso)*, *Bomanins (Bom)*, *Metchnikowin (Mtk)*, *Turandots (Tots)* and *puckered (puc)*. *Jra*, the transcription factor downstream of JNK signalling, is also upregulated. The corresponding immune pathways for each gene are marked (Y-axis) in Fig. 2D. In the ‘Disease’ context, many defence genes were found to be upregulated in *VAP^P58S^* at day 15 and day 20. Transcripts with higher values include *Tots, Atts*, *Cecs* and *Boms* amongst others. Interestingly, the peptidoglycan recognition proteins are mildly upregulated, while *pirk*, a negative regulator of immune signalling, is downregulated (Fig. 2D). We selected a few AMPs and, in an independent experiment, performed a qRT-PCR on *Drosophila* heads. Data for 5- and 20-day-old *VAP^WT^*and *VAP^P58S^* flies are shown in Fig. 2E and 2F, respectively. The results were comparable to those from 3’ mRNA sequencing, with 5-day showing no change in AMP levels and 20-day showing significantly elevated AMP levels. In agreement with the male data, *VAP^P58S^* females also showed low-grade increase in immune-responsive gene transcripts, with age. GO analysis showed terms like ‘regulation of antifungal innate immune response’, ‘innate immune response’ and ‘humoral immune response’ amongst others (Suppl. Fig. 3B). A heatmap showing the significantly DEGs showed elevated levels of defence genes (Suppl. Fig. 3C), in agreement with the male transcriptome. As observed for the male data (Fig. 2), the genes were mapped to ‘Ageing’ and ‘Disease’ (Suppl. Fig. 3D). *AMP* transcripts across Toll, IMD and Jak-STAT pathways showed a weak upswing in transcripts with age, when compared to *VAP^WT^*. When the male and female datasets were compared, females showed higher log_2_FC values overall, especially for ageing. qRT-PCR data for 5-day and 20-day (Suppl. Fig. 2E and 2F), respectively, for a few transcripts confirm the trend in the transcriptomics data.

### *VAP^P58S^* flies show age-dependent changes in neuronal function

Since *VAP^P58S^* animals show age-dependent motor deficits, we also examined changes in transcripts for genes involved in neuronal function (Fig. 3; Suppl. Fig. 4 & 5). Gene Ontology (GO) analysis of the DEGs for *VAP^P58S^*(20 and 15 day, normalized to day 5) suggests that, in addition to upregulation of immune response genes, genes involved in categories such as ‘Synaptic Transmission’, ‘Membrane potential’, ‘inorganic ion transport’ are downregulated in both male and female *VAP^P58S^* flies with age (Fig. 3A and 3B; Suppl. Fig. 5A and 5B, respectively). The genes were classified as ‘Neurotransmitter receptors’, ‘Transporters and ion channels’, ‘Synaptic Vesicle release machinery’, Synapse formation and plasticity’ and ‘Synaptic Signalling and modulators’. log2FC values were tabulated for both *VAP^P58S^* and *VAP^WT^* (Fig. 3C). Though the overall trend is for a mild decrease in transcripts with age, the reduction is more substantial in *VAP^P58S^* and affects a larger subset of genes. This suggests an overall acceleration of the decline in levels of *VAP^P58S^* neuronal transcripts in comparison to *VAP^WT^*, indicating enhanced ageing. In accordance, *VAP^P58S^* females also showed downregulation of most neuronal genes with age compared to *VAP^WT^* (Suppl. Fig. 5C). Heatmaps and volcano plots for males and females are shown in Suppl. Fig. 4A, 4B and Suppl. Fig. 5D, 5E, respectively.

**Figure 3:**
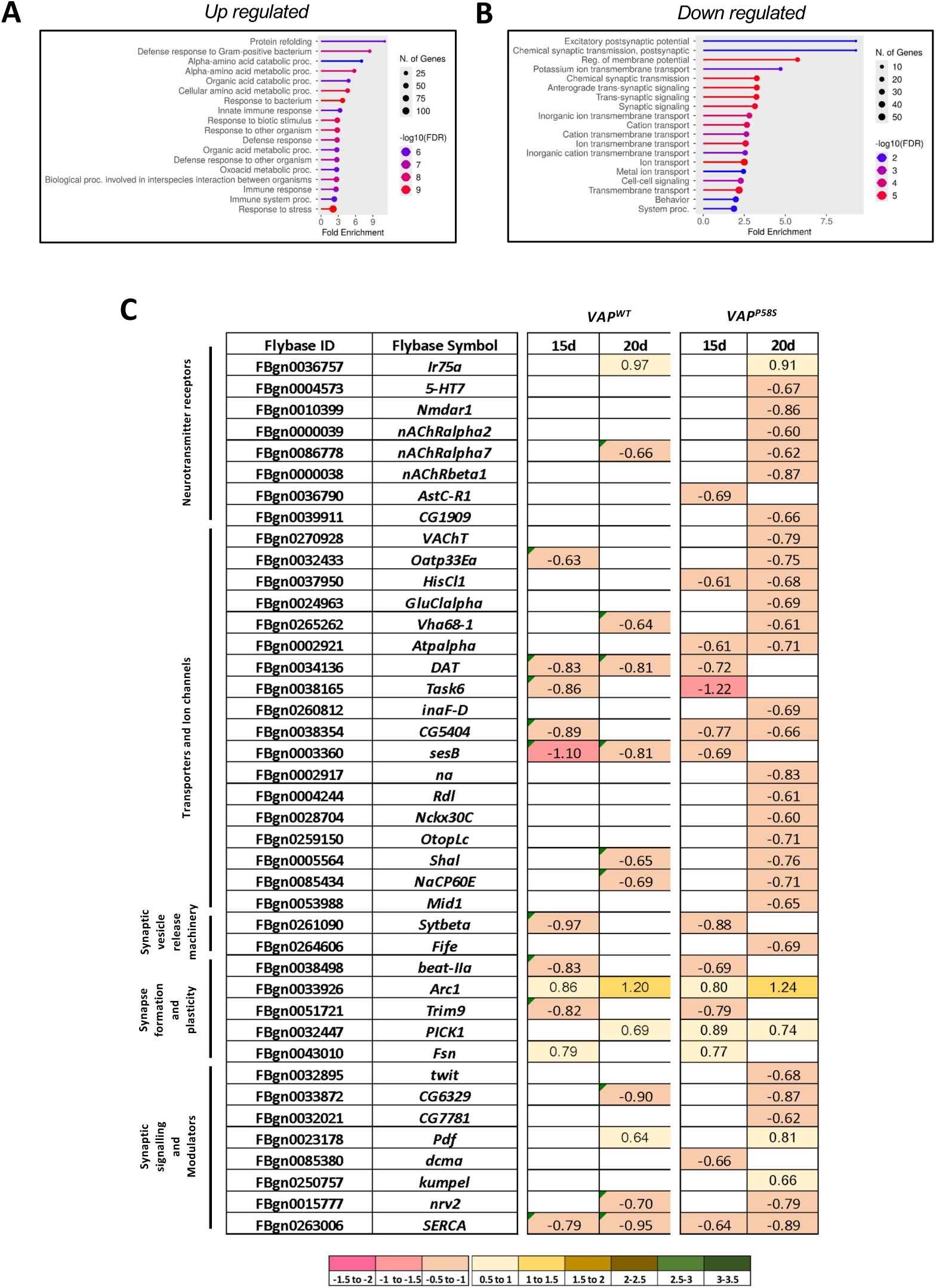
Transcriptomic analysis showing age-dependent changes in synaptic signalling in *VAP^P58S^* males. **A.** GO analysis for significantly upregulated genes in *VAP^P58S^* between day 5 and days 15/20. Suppl. Fig. 4 displays the heat map and volcano plots for these datasets, while Suppl. Fig. 5 displays data for females. **B.** GO analysis for significantly downregulated genes in *VAP^P58S^* between day 5 and days 15/20. **C.** Tabular representation of log_2_FC values for synaptic gene transcripts. The genes have been categorized based on the functions they perform. The table shows comparison of *VAP^P58S^* at days 15 and day 20 with *VAP^P58S^* at day 5. The same comparison has been done for *VAP^WT^*.

Our observations in the *VAP^P58S^* transcriptome align well with the findings from multiple independent head transcriptomic studies that evaluated ageing and age-related disorders in *Drosophila*. These transcriptomes show a downregulation of transcripts associated with synaptic transmission at various levels. The neural gene categories listed above have been repeatedly enriched in these studies. Broad classes of neurotransmitter receptors like cholinergic nAChR receptors, glutamate receptors and NMDAR have been shown to be downregulated with age. This decline is consistent with the age-associated dampening of synaptic responsiveness. Additionally, a downregulation in genes involved in regulating excitability like Na^+^/K^+^/Cl^−^ ion channels such as such as *Na^+^ pump α-subunit (Atpalpha)* and *stress-sensitive B (sesB)* in these brain transcriptomes points towards impaired electrical signalling (Pacifico et al., 2018). Moreover, downregulation of *Shaker cognate I (ShaI)* protein levels is known to be associated with age-dependent motor decline in flies (Vallejos et al., 2021). Apart from these, components of the synaptic vesicle formation and release machinery including *Synaptotagmins (Syts)* show decreased expression with age reflecting weakening of the synapse and loss of synaptic plasticity (Pacifico et al., 2018).

Neuropeptides like *Pigment-dispersing factor (Pdf)*, which is involved in maintaining normal circadian rhythm, similarly show an increase in expression, indicating changes in broader circuit neuromodulation (Umezaki et al., 2012). Overall, the coordinated downregulation of these receptors, ion channels, transporters, along with vesicle release, and structural synapse-associated genes, points towards disruption of the synaptic structure as well as function as flies age. Additionally, the ageing transcriptomes in these various studies show increased expression of immune-associated genes.

Our transcriptomic analysis of *VAP^P58S^* flies thus shows age-associated immune dysregulation and synaptic dysfunction in both male and female *VAP^P58S^* mutants. This further underscores inflammation as a key pathological mechanism in disease progression. Additionally, our study supports the idea that the *VAP^P58S^*flies undergo accelerated or premature ageing marked by loss of neuronal gene expression and heightened proteostasis stress.

### *VAP^P58S^* gut does not show inflammatory changes

In addition to our transcriptomic experiments on the heads of the *VAP^P58S^* mutants, we performed age-dependent transcriptomics on the gut. Our goal was to uncover significant transcriptomic changes in the gut of *VAP^P58S^*, both in the context of the gut:brain axis and effects of *VAP^P58S^* on ageing and disease within the gut. Unlike in the brain, the gut does not show large changes in gene expression either in the ‘ageing’ or the ‘disease’ context (Suppl. Fig 6). A volcano plot for the gut DEGs (Suppl. Fig. 6A) between *VAP^P58S^* and *VAP^WT^,* highlights the few genes that show change. GO analysis highlights that these genes are involved in disaccharide metabolic process, oogenesis, sexual reproduction and cell development. *VAP^P58S^* flies show reduced egg lay (unpublished data) and thus may have reproductive defects, which would explain the GO terms relating to reproduction being identified. Unlike in the head transcriptome, we could not identify trends that associated with defence response or immunity (Suppl. Fig. 6B). A heatmap of the DEGs between *VAP^P58S^* and *VAP^WT^* at 5-day and 20-day (Suppl. Fig. 6C) shows a set of genes up- and downregulated, with very few immune genes identified (Suppl. Fig. 6D). These changes do not lead us to any leads towards a significant role for the gut:brain axis in the *VAP^P58S^* model. The experiments were first carried out in females, and because the *VAP^P58S^* gut did not show significant changes, we did not generate data for the male gut transcriptome.

In conclusion, data from the gut transcriptomics of *VAP^P58S^* shows minimal changes, suggesting that neuroinflammation in the *VAP^P58S^* adult brain is not correlated with changes in the gut transcriptome.

### A glial enhancer/suppressor screen identified Kay as a key regulator of motor function in *VAP^P58S^* flies

Previous work from our lab using the *ΔVAP;gVAP^P58S^* model identified the IMD pathway as a regulator of glial inflammation (Tendulkar et al., 2022). In this study, we screened genes in the Toll and the JNK pathways to establish their roles in the ALS8 context*. VAP^P58S^; RepoGal4* females were crossed to either knockdown (KD), overexpression (OE), dominant negative (DN) or constitutive active (CA) *UAS* lines of *dorsal* (*dl*), *dorsal-like immune factor* (*dif*), *cactus* (*cact*), *spatzle* (*spz*), *spatzle 2* (*spz2*), *spatzle 5* (*spz5*), *spatzle 6* (*spz6*) and *Toll10b* for the Toll pathway or *kay*, *jun-related antigen* (*jra*), *basket* (*bsk*) and *hemipterous* (*hep*) for the JNK pathway. F1 males were selected with climbing ability being used as the readout (Suppl. Fig. 7A). Gene modulations that worsened the climbing ability of the *VAP^P58S^*flies were termed as ‘Enhancers’, whereas those that improved the climbing ability were considered as ‘Suppressors’ (Suppl. Fig. 7B).

The first pathway we examined was the Toll pathway. Suppl. Fig. 7C depicts climbing assays performed on days 5, 10, and 15. KD of *dl* (p>0.05, 2, light blue), *dif* (L1) (p>0.05, 3, light green), *dif* (L2) (p>0.05, 4, dark green), *cact* (p>0.05, 5, pink), *spz2* (p>0.05, 8, yellow curve), *spz5* (p>0.05, 9, orange curve), *spz6* (p>0.05, 10, dark blue curve) did not show any significant change in the climbing index when compared to the age matched control *VAP^P58S^; RepoGal4* (1, black curve). OE of the active form of the classical Toll receptor, *Toll10b*, similarly showed no change (p>0.05, 11, grey curve). *cact* OE on the other hand showed mild suppression of motor defects at day 10 (p<0.05, 6, purple curve). Similarly, *spz* KD mildly suppressed motor defects at day 10 (p<0.05, 7, brown curve). Suppl. Fig. 7D is a tabulated summary with color-coded p-values. Shades from light pink to magenta show increasing levels of motor deterioration (Enhancers), whereas shades from yellow to dark green depict improvement in motor function (Suppressors).

The second pathway that we examined was the JNK pathway. Suppl. Fig. 7E shows climbing assays for the modulation of JNK pathway genes. KD of *kay* showed significant enhancement of the motor function defect in the *VAP^P58S^* flies at day 15 (p<0.001, 2, red curve) whereas *kay* OE improved climbing ability at day 10 (p<0.001, 4, light green curve). OE of *Fra.Fbz* (L1), the dominant negative form of *kay* showed no change in climbing index (p>0.05, 3, dark pink), while OE of *Fra.Fbz* (L2) was lethal. OE of *jra^DN^* (p<0.01, 7, blue curve) as well as KD of *bsk* (p<0.05, 8, sea green curve) showed mild enhancement of motor defects at day 15. OE of a *bsk^DN^*, on the contrary, showed very mild suppression (p<0.05, 9, orange curve). *jra* KD (p>0.05, 5, ochre curve) and OE of either *jra* (p>0.05, 6, pink curve), *bsk* (p>0.05, 10, purple curve) or *hep* (p>0.05, 11, grey curve) did not affect the climbing index. OE of the constitutively active *hep* was lethal. A summary table presents the p-values for different intervals, color-coded as previously described (Suppl. Fig. 7F).

Our glial screen thus indicates that the JNK pathway is involved in regulating motor function in *VAP^P58S^*, and we have focused subsequent experiments on understanding *kay’s* glial role in the disease model in greater detail.

### *kay*/*kay^SCR^* OE in glia partially rescues motor function

Because the glial enhancer/suppressor screen identified *kay* as a critical modulator of motor function in *VAP^P58S^* flies, we generated additional *kay* lines to validate the rescue. Of the two variants, one expresses wildtype *kay* (*UAS-kay*) and the second expresses the small ubiquitin-like modifier (SUMO) conjugation resistant (SCR) form of *kay* (*UAS-kay^K357R^*) (further referred to as *UAS-kay^SCR^*). In mammals, SCR Fos is resistant to SUMO conjugation and is a more active form (Bossis et al., 2005). Hence, we included it in our OE experiments as an alternate allele with the expectation that it would give a similar or a stronger rescue of motor function.

Fig. 4A shows the climbing index curve for glial modulation of *kay* in the background of *VAP^P58S^*, with Fig. 4C tabulating enhancement/suppression of *VAP^P58S^* function based on p-value. KD of *kay* in the glia, showed significant worsening of motor function defects in the *VAP^P58S^* flies by day 15 (p<0.01, 2, red curve) as compared to *VAP^P58S^; RepoGal4/+; +* control (1, black curve). OE of *kay*, with the line procured from BDSC (L1) showed significant rescue of motor function at day 10 (p<0.0001, 3, green curve). OE *kay* (L2) using the laboratory generated line as well as *kay^SCR^* line showed a stronger rescue of motor function at day 10 and day 15 (p<0.0001, 4 and 5, light blue and dark blue curves, respectively).

**Figure 4:**
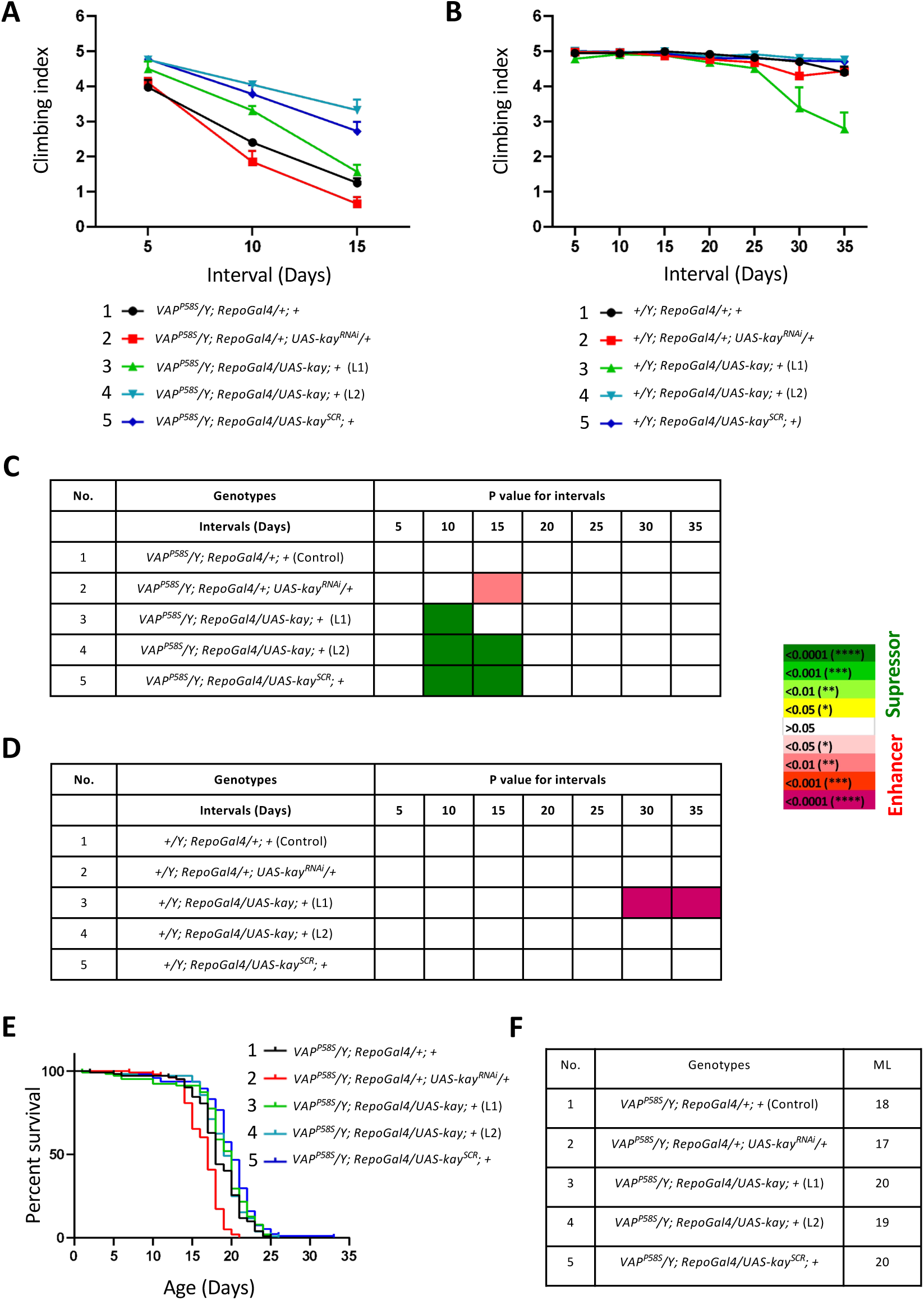
Glial OE of *kay* or *kay^SCR^* partially rescues motor function and lifespan of the disease model. **A.** Climbing indices for glial KD of *kay* (2, red), *kay* OE (L1) (3, light green), *kay* OE (L1) (4, light blue), *kay^SCR^* (5, dark blue) in the background of *VAP^P58S^*. *VAP^P58S^/Y; RepoGal4/+; +* was used as control (1, black). Values shown as mean ± SEM. n=25-30, N=3 **B.** Climbing indices for glial KD of *kay* (2, red), *kay* OE (L1) (3, light green), *kay* OE (L1) (4, light blue), *kay^SCR^* (5, dark blue) in the background of wildtype. *+/Y; RepoGal4/+; +* was used as control (1, black). Values depicted as mean ± SEM. n=25-30, N=3 **C and D.** Summary table for the gene modulations in figures (A) and (B), respectively. The individual P values have been colour-coded. Statistical analysis as previously described. **E.** Lifespan curves for glial KD of *kay* (2, red), *kay* OE (L1) (3, light green), *kay* OE (L1) (4, light blue), *kay^SCR^*(5, dark blue) in the background of *VAP^P58S^*. *VAP^P58S^/Y; RepoGal4/+; +* was used as control (1, black). The combined p-value for the set is p<0.0001. n=80-100, N=3 **F.** Results tabulated in terms of median lifespan for each genotype.

**Figure 5:**
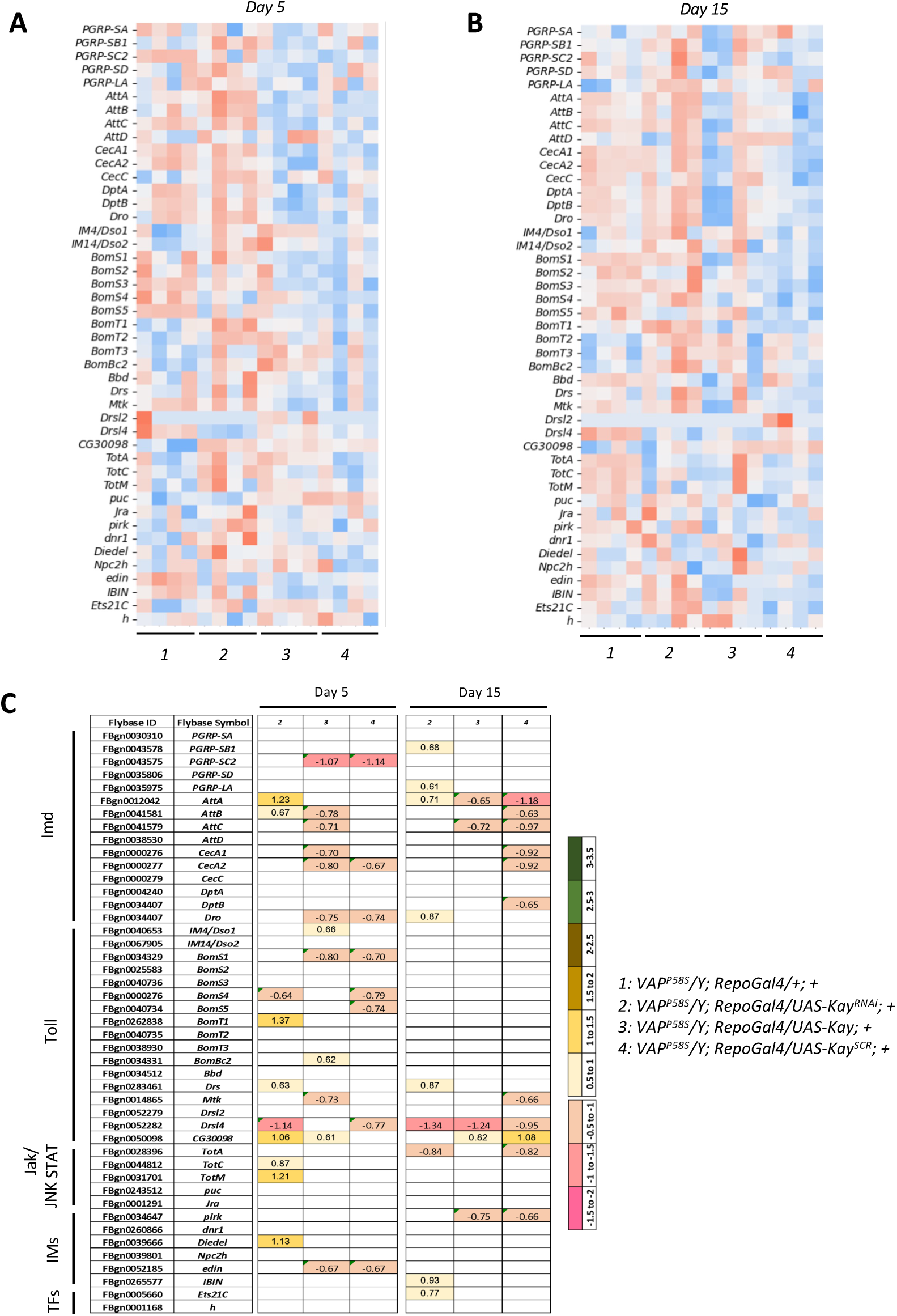
*kay* or *kay^SCR^* in glia rescues neuroinflammation in the *VAP^P58S^* flies. **A.** Heatmap showing the normalized expression counts for immune genes significantly differentially expressed in 2, *VAP^P58S^/Y; RepoGal4/+;UAS-Kay^RNAi^*; 3, *VAP^P58S^/Y; RepoGal4/UAS-Kay;* 4, *VAP^P58S^/Y; RepoGal4/UAS-Kay^SCR^* with 1, *VAP^P58S^/Y; RepoGal4/+* being used as the control at day 5. **B.** Heatmap showing the normalized expression counts for immune genes significantly differentially expressed in 2, *VAP^P58S^/Y; RepoGal4/+;UAS-Kay^RNAi^*; 3, *VAP^P58S^/Y; RepoGal4/UAS-Kay;* 4, *VAP^P58S^/Y; RepoGal4/UAS-Kay^SCR^* with 1, *VAP^P58S^/Y; RepoGal4/+* being used as the control at day 15. **C.** Log_2_FC values plotted for significantly differentially expressed immune genes for day 5 and day 15. The colour code and genotypes have been mentioned on the side.

Next, we looked at the effect of all the modulations on motor function in wildtype flies (Fig. 4B, 4D). Flies overexpressing *kay* showed deterioration of motor function day 30 onwards (p<0.0001, 3, green curve) which was in contrast to the rescue of motor function observed in the *VAP^P58S^* flies. *+/Y; RepoGal4/+; +* was used as the control (1, black curve).

The *VAP^P58S^* flies exhibit reduced lifespan, in addition to motor defects. As *kay* KD worsened motor function in these flies and *kay/kay^SCR^* rescued it, we examined if lifespan showed a similar trend. *VAP^P58S^/Y; RepoGal4/+; +,* which was used as the control, showed an ML of 18 days (Fig. 4E and 4F, 1, black curve). KD of *kay* showed a very mild reduction of lifespan with the ML being 17 days (Fig. 4E and 4F, 2, red curve). OE of *kay* (L1), *kay* (L2) as well as *kay^SCR^* showed a mild increase in the lifespan with MLs being 20, 19 and 20 days (Fig. 4E and 4F, 3, 4 and 5, green, light blue and dark blue curves), respectively.

Thus, our experiments establish *kay* as a modifier of the disease with OE in glia, leading to an improvement in the motor deficit of *VAP^P58S^* males.

### Glia *kay/kay^SCR^* OE mitigates age-dependent neuroinflammation

*VAP^P58S^* flies show age-dependent neuroinflammation in the head. Further, glial knockdown of *kay* exacerbated age-dependent motor deterioration in *VAP^P58S^* mutants, whereas its OE suppressed these motor defects. This would suggest that motor function in the *VAP^P58S^* model was a function of the extent of inflammation. We therefore OE & KD *kay* in the head and examined changes in the progressive low-grade inflammation observed in the disease model.

Fig. 5 displays the set of immune genes identified as upregulated in *VAP^P58S^* mutant males, many of these defense genes, visualised using their normalised expression profiles across four genotypes (1, *VAP^P58S^/Y; RepoGal4/+;* 2, *VAP^P58S^/Y; RepoGal4/+;UAS-Kay^RNAi^*; 3, *VAP^P58S^/Y; RepoGal4/UAS-Kay;* 4, *VAP^P58S^/Y; RepoGal4/UAS-Kay^SCR^*) and two timepoints. The heatmaps for both day-5 (Fig. 5A) and day-20 (Fig. 5B) indicate differences in immune gene expression across the four genotypes. Upon *kay* KD, the expression of immune genes was elevated compared to age-matched controls. In contrast, overexpressing *kay* or *kay^SCR^* reduced immune gene expression at both 5 and 20 days (Fig. 5A and 5B, respectively). Intriguingly, the reduction in immune gene expression obtained with overexpression of *kay^SCR^* was better or at par with that seen with *kay^WT^*. This outcome reinforces previous conclusions from mammalian studies that SCR Fos is an active form (in this case, a stronger inhibitor of immune gene expression). To quantify gene changes, we plotted the log_2_FC values for these immune genes. Most AMPs from Imd, Toll as well as Jak/STAT pathways were found to be elevated upon *kay* KD in the glia at both timepoints. Glial overexpression of *kay* or *kay^SCR^* on the other hand downregulated the expression of the AMPs. The strongest reduction was seen in *AttA*. Additionally, *PGRP-SC2* also showed a significant downregulation of expression. Other AMPs like *AttB*, *AttC*, *CecA1*, *CecA2*, *DptB*, *BomS4*, *BomS5*, *Mtk* and *TotA* also showed milder reduction in expression (Fig. 5C).

Is Kay a known regulator of defence genes, and does the literature indicate a potential mechanism for Kay-modulated transcriptional repression? *kay* was one of the 45 genes (Fisher et al., 2023) modulated in the embryo to study their transcriptional targets during embryogenesis (10-12 hours and also 16-18 hours post-fertilisation). Fisher *et al*. 2023 found that *kay* primarily regulated 25 defence response genes, including Bomanin’s, showing ∼30% overlap with defence genes identified in our study (Suppl. Fig 8A, 8B and 8C). Importantly, all these genes were negatively regulated by Kay, with *kay* KD leading to their upregulation. Further, we analyzed Chromatin Immunoprecipitation (ChIP)-sequencing (Seq) data from the Encyclopedia of Regulatory Networks (modERN) consortium and mapped Kay occupancy to the gene bodies of all genes in *Drosophila.* We found enriched peaks for Kay binding on the promoters of 4762 genes (Suppl. Fig. 8D, Clusters 2,4 and 5). However, the Kay responsive genes identified in our study did not appear to have binding sites for Kay, suggesting that Kay acts indirectly, possibly regulating transcription via another transcription factor. In the few cases where Kay showed enriched binding to promoters of genes, such as *rel*, *dl* and *STAT92E*, our mRNA sequencing data showed no change in the expression of these genes upon *kay* modulation (Suppl. Fig. 8E).

Our transcriptomic data thus suggest that OE of *kay* or *kay^SCR^*contributes to a significant rescue of climbing ability in the *VAP^P58S^* mutant by clearing neuroinflammation. We believe this is achieved via Kay’s broad repression of major immune pathways. On the other hand, the absence of promoter binding and the lack of changes in the expression levels of the major immune TFs suggest an indirect mechanism of regulation by Kay.

## Discussion

Neuroinflammation is a pathological hallmark of most neurodegenerative disorders, including ALS (Johnson and Lukens, 2025; Thonhoff et al., 2018; Zhang et al., 2023; Zhao et al., 2013). Though ALS was initially considered a disease primarily associated with motor neuron degeneration, the involvement of non-neuronal cells like glia is being increasingly explored and established (Boillee et al., 2006). Mounting evidence points to a glia-derived pro-inflammatory state in various animal models (Koehn et al., 2025; O’Rourke et al., 2016; Yamanaka et al., 2008) and in human ALS patients (Femiano et al., 2024; Gagliardi et al., 2024; Mishra et al., 2017; Steinacker et al., 2018).

Glial neuroinflammation has emerged as a central non-cell-autonomous mechanism in ALS. In neurodegeneration, microglia shift to a chronically activated, pro-inflammatory phenotype driven by signals such as misfolded proteins, damage-associated molecular patterns (DAMPs), and cytokines (Chiu et al., 2008; Weydt et al., 2004) (Beers et al., 2006; Martens et al., 2022). Activated microglia release Tumour Necrosis Factor TNF-α, Interleukin (IL)-1β, IL-6, ROS, and nitric oxide, and activate the NOD-like receptor pyrin domain-containing protein 3 (NLRP3) inflammasome, amplifying local inflammation and promoting motor-neuron vulnerability (Deora et al., 2020; Nagai et al., 2007; Xu et al., 2025). These reactive astrocytes, termed ‘A1 astrocytes’ (Liddelow et al., 2017), reduce metabolic and synaptic support to neurons. Further, peripheral immune cells infiltrate the CNS, compromising the blood-brain barrier (Beers et al., 2008; Henkel et al., 2004). Neurons, glia and secreted pro-inflammatory factors create a chronic immune response that accelerates motor-neuron degeneration and disease progression in ALS. *Drosophila* models of ALS also support the central role of glial ‘activation’, their phagocytic roles, demonstrating activation of Toll/IMD or JNK pathways in the brain in neurodegeneration or traumatic brain injury models (Cao et al., 2013; Elguero et al., 2023; Katzenberger et al., 2013; Kounatidis et al., 2017; Lee et al., 2020; Li et al., 2018; Tendulkar et al., 2022; van Alphen et al., 2022).

In our *Drosophila VAP^ALS^* model, we observe an increase in low-grade inflammation as animals age and lose motor function. The progressive decline in motor function correlates with increasing inflammation. A targeted enhancer/suppressor motor function screen identified the JNK transcription factor Kay as a key disease modifier. Kay, along with its partner Jra, forms a heterodimeric Activator Protein (AP)-1 complex (Kockel et al., 2001; Perkins et al., 1990) implicated in diverse physiological pathways, including stress response (Alfonso-Gonzalez and Riesgo-Escovar, 2018; Biteau et al., 2011; Kockel et al., 2001; Perkins et al., 1990; Tafesh-Edwards and Eleftherianos, 2020). The AP-1 complex (Fos/Jun in vertebrates) is assumed to be the most stable form, though both Kay and Jra can function as homodimers (Alfonso-Gonzalez and Riesgo-Escovar, 2018). JNK signalling, with Kay/Jra as transcriptional regulators, is a major regulator of the immune response (Kleino and Silverman, 2014; Park et al., 2004; Silverman et al., 2003; Tafesh-Edwards and Eleftherianos, 2020). Both Jra and Kay are expressed in neurons and glia and their vertebrate orthologs have been implicated in neurodegenerative diseases (Gogia et al., 2020; Sahana et al., 2023).

We find that KD of *kay* in the glia led to a worsening of motor function, while OE of *kay* or *kay^SCR^* improved motor function in *VAP^P58S^*mutant lines. Kay appears to function as a broad regulator of the basal immune response, affecting defence genes downstream of multiple immune pathways. Since Kay is a transcriptional regulator, and is negatively regulating defence genes, one possible mechanism is for it to bind to promoters of defense genes and function as a dampening factor, however our analysis appears to rule this out as a primary mechanism.

What then is the link between VAP, an ER-associated protein with Kay? How does this relationship, in glial cells, regulate age-dependent progression of gene-regulatory networks, especially those related to innate immune pathways? How does a point mutation in VAP lead to accelerated ageing and inflammation? A hint comes from studies in viral immunity (Ji et al., 2025), which links VAPB in mammals to innate immune signalling via the cyclic guanosine 5′-monophosphate (GMP)–adenosine 5′- monophosphate (AMP) synthase (cGAS) - Stimulator of Interferon genes (STING) pathway (Decout et al., 2021; Dvorkin et al., 2024). The cGAS-STING pathway responds to aberrant cytosolic DNA and is an evolutionarily conserved mechanism for defence against microorganisms (Marques et al., 2024). (Ji et al., 2025) find that VAPB binds to and sequesters the ER-resident STING under basal conditions. In response to cytoplasmic dsDNA stimulation, STING translocates away from the ER, presumably to the Golgi or peroxisomes, and this perinuclear translocation triggers an inflammatory response. The authors implicate TANK-binding kinase 1 (TBK1) as a central facilitator of the VAPB/cGAS-STING pathway, which phosphorylates and activates downstream transcriptional regulators. VAPB thus acts as a negative regulator of the STING-associated immune response (Ji et al., 2025). TBK1 is one of a host of cytoplasmic kinases (CKs) that function as crucial hubs in signaling, acting as molecular switches that phosphorylate targets to regulate transcriptional activation and repression. The CKs integrate signals, forming complexes with receptors, and are essential for both initiating and fine-tuning innate and adaptive immunity. The CKs are associated with cellular compartments, including the plasma membrane, the Golgi, endosomes, and the ER. CKs can also travel to the nucleus as part of their activation. In *Drosophila,* CKs such as Pelle, Hopscotch and TAK1 are upstream of the major immune pathways and fine-tune the expression of defence genes. The IκB kinase epsilon (IKKε) is the fly ortholog of TBK1 and functions to regulate the degradation of Death-associated protein inhibitor of apoptosis 1 (DIAP1), a central regulator of apoptotic signalling. STING is also conserved in flies (Liegeois and Ferrandon, 2022), with (Martin et al., 2018) demonstrating a role for *Drosophila melanogaster* STING (dmSTING) in the immune response. (Martin et al., 2018) find that dmSTING associates with cyclic dinucleotides and utilises the IMD pathway to activate defence genes. Intriguingly, TGF-β-activated kinase 1(TAK1), a MAP3K kinase related to TBK1, is a key regulator of NFκB signalling in *Drosophila*, and is upstream of both REL (Silverman et al., 2003; Vidal et al., 2001) and JNK (Geuking et al., 2005; Igaki et al., 2002) pathways.

In our model (Fig. 6), we propose that in *Drosophila*, VAP regulates Toll, JAK-STAT JNK and IMD signalling via its action on upstream CKs. In the *VAP^P58S^* mutant, ER structure, function, and dynamics are affected, leading to CK modulation. Multiple VAP interactors, as exemplified by STING, may also directly influence CK function. Specifically, we suggest that TAK1 function decreases with age in *VAP^P58S^* animals. This leads to reduced basal phosphorylation of Kay/Jra, which, in turn, depresses defence genes (Fig. 6). The low-grade inflammation seen with increasing age would be a combined result of TAK1 modulation of its two arms – the ‘activating’ REL arm vs the ‘negative-regulating’ Jra/Kay arm. The seat of the basal increase in inflammation is the glia, and sustained chronic low-grade inflammation leads to non-cell-autonomous spread to neurons, which in turn leads to neuronal degeneration.

**Figure 6:**
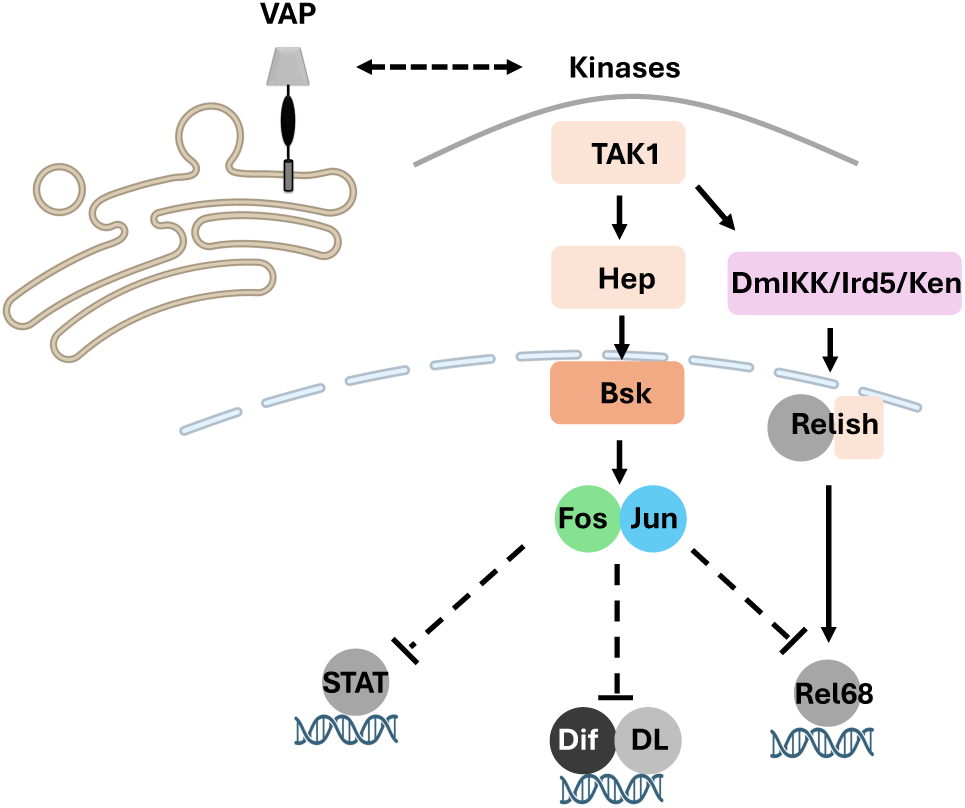
Model for regulation of the glial immune response. *VAP^P58S^* flies show age-dependent low-grade enhancement of neuroinflammation, evident from the increased expression of defence genes downstream of the Imd, Toll, JNK and JAK-STAT pathways. We propose that either a Kay homodimer or a Jra/Kay heterodimer negatively regulates defence genes by repressing transcriptional activation of target genes in glia. In *VAPB^P58S^,* this negative regulation is weakened with age, allowing for a progressive increase in inflammation. The increase in inflammation is correlated with an age-dependent motor deficit. Glia inflammation, in turn, is predicted to lead to the secretion of inflammatory factors that affect glial: neuronal homeostasis and ultimately neuronal cell death.

The discovery of the VAPB:STING interaction (Ji et al., 2025) also highlights the possibility of STING being a significant mechanistic pathway for the initiation and progression of ALS (Almalki et al., 2025; Decout et al., 2021; Ji et al., 2025), with the source of the triggering DNA being damaged mitochondria in glial cells. This possibility, for the *VAP^P58S^* model, will be tested in future experiments in our laboratory.

In summary, we find that Kay is a novel neuroinflammatory glial modulator in the *VAP^P58S^* model, suggesting an equivalent role for Fos in human microglia. Although the mechanism of this regulation or the link between VAP and Kay remains unclear, as is Kay’s ability to regulate a broad set of inflammatory genes, further targeted mechanistic studies can open avenues for the development of new treatment strategies for ALS.

## Materials and Methods

### Fly husbandry

All flies were reared at 25 ⁰C on standard cornmeal agar in a 12-hour light-dark cycle. In specific cases, crosses were placed at 18 ⁰C to reduce developmental effects, then flies were collected on the first day of eclosion and transferred to vials at 25 ⁰C. All subsequent experiments were performed at 25 ⁰C. The fly lines used in this study are tabulated in Suppl. Table 1.

### Generating the *VAP^P58S^* mutant using CRISPR-Cas9 genome editing

We employed a CRISPR-Cas9 ssODN-based strategy to generate a mutation at the *VAP* locus. In this strategy, a Double Strand Break (DSB) at the target locus is created by a guide RNA (*gRNA*), followed by repair of the DSB using a single-stranded oligodeoxynucleotide (*ssODN*) template containing the mutation. To generate this DSB at the *VAP* locus, as a first step we designed and cloned a *VAP* gRNA containing the sequence 5’-TACGCAGTAGCGTTTCG-3’ (Suppl. Fig. 1A) in the pBFv-U6.2B vector (Kondo and Ueda, 2013). The gRNA sequence targeted 3 bp upstream of the planned mutation site. The *pBFv-U6.2B VAP-gRNA* plasmid was injected into the *v^−^;attP40* line, and transgenics were scored by screening for *v^+^*. The stable balanced homozygous *VAP-gRNA* line was then crossed to a *v^−^*; *nanos-Cas9* (*nos-Cas9*) line (Kondo and Ueda, 2013) (Suppl. Fig. 1B). Young embryos (0-2 hours) of the F1 generation were collected. Approximately 700 embryos were injected with a synthesized ssODN (*5’-TTGATCGAAAGGGAATTATCTTGCCGATGTTGCTACGTACGCAGTAGCGTTTCGGCGCGGTTGTCTTGATCTTGAAGACCAGAGGCAGAGCCGAGTTGTTG-3’*) that contained the mutant sequence (Suppl. Fig 1D). Once the embryos developed into adult flies, straight-winged flies were crossed to balancers to preserve the genetic mutation and then stabilised, before screening for the mutation by genomic PCR. The ssODNA design incorporated a restriction digestion site 5’TACGTA-3, digested by SnaBI (Suppl. Fig. 1C). Of the 359 lines stabilized, the incorporation of the template appeared efficient, as 8 of the first set of 50 lines screened showed the presence of a SnaBI site (Suppl. Fig. 1C). Four *VAP^P58S^*and three *VAP^WT^* lines, validated by sequencing were maintained, while all remaining lines were discarded. These lines did not contain the gRNA (*v^−^*, *VAP^P58S^*) and were cleaned by crossing to a *v^−^* mutant line and rebalancing with FM7a.

### RNA isolation, 3’mRNA library preparation and sequencing

Total RNA was isolated from either 50 heads or 50 guts using trizol-chloroform extraction. For each sample, 3’ mRNA libraries were generated from four biological replicates using the QuantSeq 3’ mRNA-Seq Library Prep Kit FWD (Lexogen, 015.96 and #191.91). Library concentrations were quantified with the Qubit™ dsDNA HS Assay Kit (Thermo Fisher Scientific, #Q32851). Quality assessment and library size estimation was done using an HS DNA kit (Agilent, #5067-4626) in a Bioanalyzer 2100 for individual libraries. Equimolar amounts of each library were pooled and single-end 75bp reads were sequenced on the Illumina novaseqx plus platform.

### Read mapping, counts generation, differential expression analysis and data visualization

Sequencing reads were aligned to the *Drosophila melanogaster* reference genome (version dm6) using the STAR aligner. Gene-level expression quantification was carried out using HTSeq-count, which generated raw read counts for each gene based on the corresponding annotation. Differential gene expression analysis was performed using DESeq2 through pairwise comparisons of the biological conditions. All computational steps, including alignment, quantification, and statistical analysis, were conducted using the Bluebee platform and Kangooroo NGS data analysis tool. Matplotlib (version 3.10.0) and seaborn (version 0.13.2) libraries were used to plot heatmaps. VolcaNoseR was used for plotting volcano plots. Downstream analysis was done using DAVID and G-profiler. GO analysis was done using ShinyGo (version 0.77).

### qRT-PCR

RNA was extracted from 40-50 heads per genotype using trizol-chloroform extraction. 1ug RNA was used to prepare cDNA using cDNA reverse transcriptase kit (Thermo Fischer Scientific, #2911843). For qRT-PCR, gene specific primers were used along with iTaq UniversalSyBr Green Supermix (Biorad, #1725124) in qTOWER^3^ Series Real-Time PCR Thermal Cycler. Normalization was done using *rp49* as the housekeeping gene. Forward (FP) and reverse (RP) primers used for the experiment are given in Suppl. Table 2.

### Cloning and construct generation for *kay/kay^SCR^* OE lines

cDNA for *kay* was amplified from the *Drosophila* gold collection library using specific primers. *kay^K357R^* was generated using site-specific mutagenesis to make a SUMO conjugation-resistant form of Kay (*Kay^K357R^/ Kay^SCR^*) by mutating lysine to arginine. The second-chromosome UAS transgenics for *Kay^WT^* and *Kay^K357R^* were generated by targeted insertion at attP40 using the pUASp-attB vector obtained from the *Drosophila* Genomics Resource Centre (DGRC, #1358). The transgenics were screened, balanced and then used for experiments. Primers used for the experiment are given in Suppl. Table 2. In addition to *UAS*-*Kay^WT^*& *UAS*-*Kay^K357^*, the BDSC:7213 (*UAS-Kay^WT^)* line from Bloomington *Drosophila* Stock Centre was used to validate and extend our results. Expression of all three lines was pupal lethal using constitutive Gal4 drivers, as is expected with *kay* OE. Levels of fold expression for the lines were measured by qRT-PCR as 17-, 12-, and 10-fold, for *UAS*-*Kay^WT^* & *UAS*-*Kay^K357^* and *BDSC UAS-Kay* respectively.

### Motor assay

Motor performance was assessed using the startle-induced negative geotaxis climbing assay. For each replicate, 30 F1 males and females were collected for each genotype and transferred to vials. Each vial contained 15 or less age-matched flies. The motor assay was performed for each genotype on days 5, 10, 15, 20 and 25. All the experiments were performed in triplicate. Each set of 30 flies was transferred to a 250mL borosilicate cylinder, and the flies were allowed to acclimatize for 1 min. The cylinder was then tapped to make all the flies settle at the bottom of the cylinder, and the timer was started. The flies were allowed to climb for 40s and the number of flies in each zone of the cylinder were counted at the end of 40s. Flies that did not climb were scored 0 (non-climbers), flies that climbed till the 80mL (7.5cm) mark were scored 3 (poor-climbers) and the flies that crossed the 80mL mark were scored 5 (good-climbers) for both males and females. The experiment was repeated three times for each set, with a 1-minute resting period between repeats. The flies were then transferred back to the vials by anesthetizing with CO_2_. To ensure that CO_2_ did not affect the climbing performance, the flies were exposed to CO_2_ only after each day’s trial. Flies were flipped every third day for males and every fourth day for females. The scores were used to calculate the climbing index (Azuma et al., 2014; Tendulkar et al., 2022). Statistical analysis was performed using Two-way ANOVA and Tuckey’s test was used to perform multiple comparisons.

### Lifespan and survival analysis

Survival assays were conducted at 25 °C. For each replicate, 80–100 F1 male and female flies per genotype were collected and distributed into vials, with no more than 15 age-matched flies per vial. The flies were flipped every third day in case of males and every fourth day in case of females to avoid death by sticking to the media. Dead flies in each vial were counted daily till the last fly of every genotype was dead. All experiments were performed in triplicate. The data was analyzed using the log-rank test. The median lifespan (ML) for each genotype was recorded.

### Statistical analysis

Experiments were performed with at least 3 biological replicates (N=3). For all QuantSeq experiments, N=4. The n (number of samples), N values, along with the statistical tests used, are listed in the figure legends for each experiment. GraphPad Prism 8.0 was used for data plotting and statistical analysis.

## Data & Resource availability

Transcriptomics data will be uploaded to a shared public server on acceptance of the manuscript.

## Contributions

NPK and GSR conceptualised the project with inputs from AS. NPK, AT, PG, AS and VK performed the experiments. NPK, GSR, PG, and AS analysed and interpreted the data. AT screened for and stabilised the genome-edited line. NPK wrote the initial draft, and GSR refined the subsequent drafts with inputs from all authors. GSR supervised the project. GSR acquired funding.

## Acknowledgements

We thank: Bloomington *Drosophila* Stock Centre (BDSC), supported by NIH grant P40OD018537, for fly stocks; Snehal Patil and Yashwant Pawar for fly media and stock maintenance; Dr Krishanpal Karmodiya for his input on the transcriptomics experiment; Sushmita Hegde and Deepti Trivedi for design of the experiment for generation of the CRISPR edited *VAP^P58S^* mutant. The fly facility at the National Centre for Biological Sciences (NCBS), Bangalore, for the injection of the *pBFv-U6.2B VAP-gRNA*, ssODNA, *UAS-kay* & *UAS-kay^SCR^*constructs in *Drosophila* embryos.

