## Supplementary material for "Fos regulates age-dependent neuroinflammation in *VAPB^ALS^*": Suppl. Files

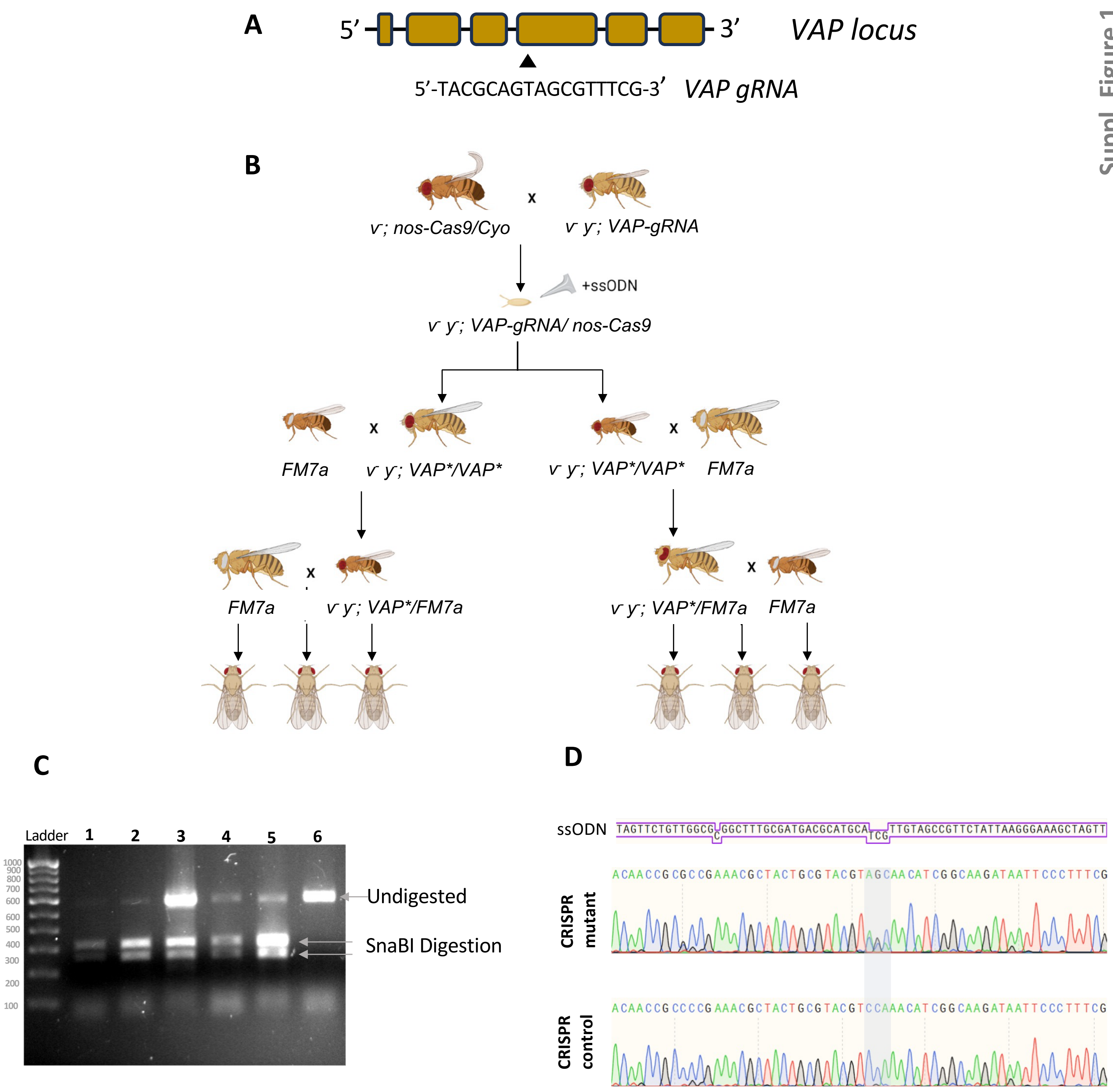


**Supplementary Figure 1: Generation of *VAP*$^{P58S}$ mutant fly lines using a CRISPR-Cas9 strategy**

**A.** Schematic for *VAP33A* genetic locus and the sequence of the *gRNA* designed to target the 4$^{th}$ exon of *VAP*.

**B.** A *VAP* guide-RNA transgenic line was generated (see Materials & Methods), balanced, and crossed to *nos-Cas9.* 700 F1 embryos were injected and 359 straight-wing flies were further balanced, expanded and stabilized.

**C.** 50 (out of 359) lines were initially screened using genomic PCR and *SnaBI* digestion. 8 lines showed digestion (e.g., Lines 2,4,5 indicating the replacement of part of the *VAP* genomic region with the mutant *ssODNA* sequence.

**D.** Eight lines were sequenced, with seven showing the replacement of codon CCA with AGC and no other mutations in the *VAP* genomic region. These were defined as 'validated' *VAP*$^{P58S}$ lines and were characterized for lifespan and startle assays. Three lines that did not show *SnaBI* digestion (panel C, lane 3, 6) were also sequenced and maintained as *VAP*$^{WT}$ (CRISPR Control) lines.

Suppl. Figure 2

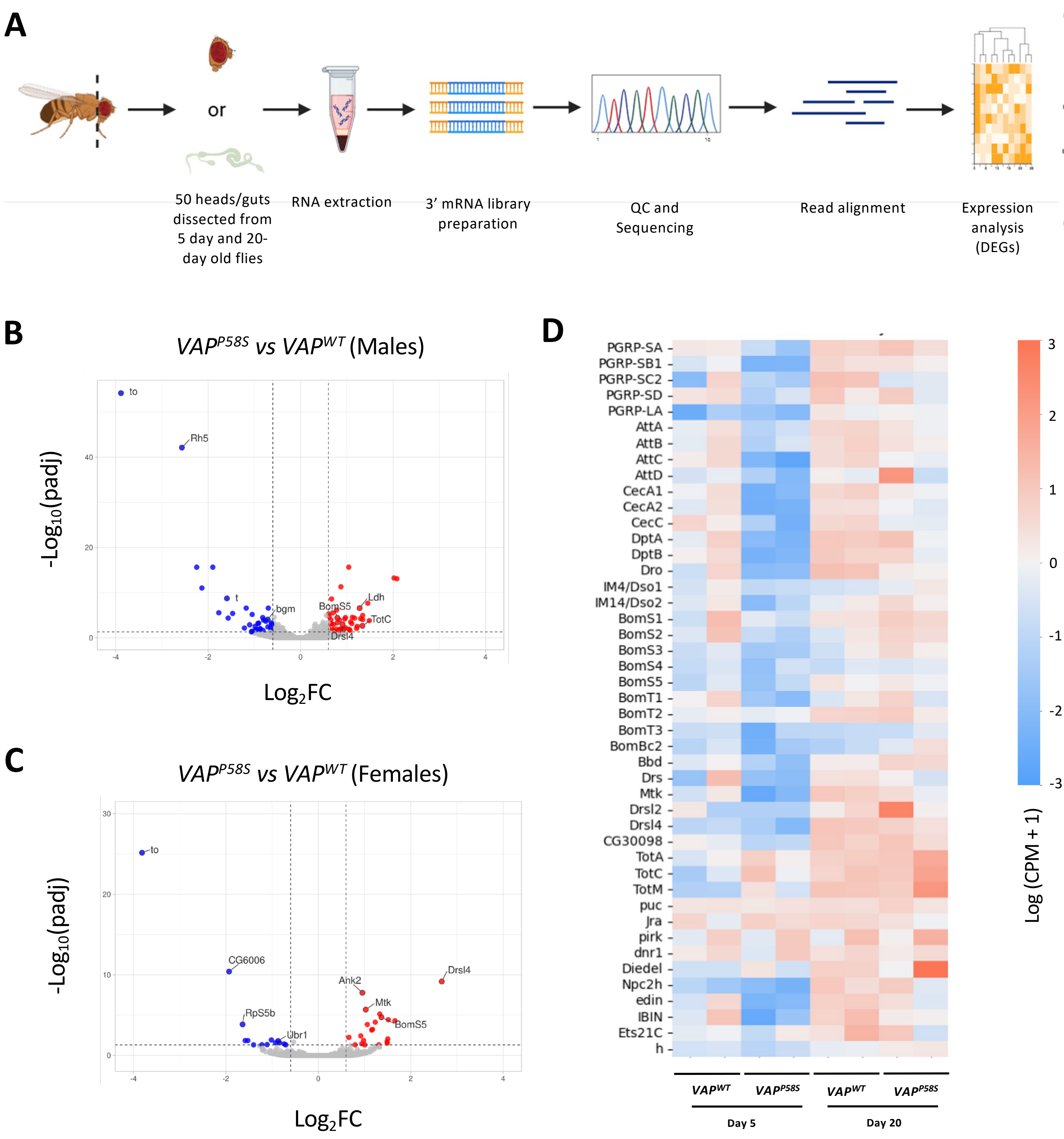


**Supplementary Figure 2: Male and female head transcriptomics for $VAP^{P58S}$ and $VAP^{WT}$**

**A.** Diagrammatic representation of the experimental workflow for 3'mRNA sequencing. 50 heads or guts per genotype per timepoint were crushed in trizol. RNA was extracted for 3'mRNA library preparation. The completed libraries were sent for quality control (QC) and sequencing following which reads were aligned and the downstream analysis to obtain DEGs was performed

**B & C.** Volcano plots for males and females respectively for the complete dataset for $VAP^{P58S}$ and $VAP^{WT}$, highlighting a few immune genes and some common genes in the two sexes

**D.** Heatmap showing the normalized expression counts for immune genes significantly differentially expressed between $VAP^{P58S}$ and $VAP^{WT}$ at day 15.

Suppl. Figure 3

A

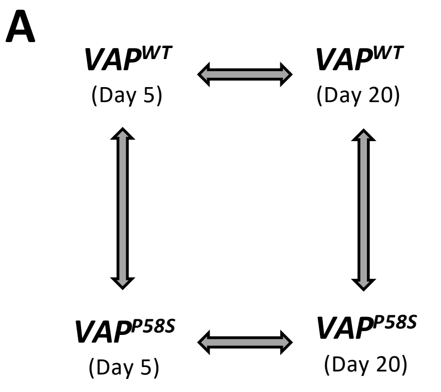


B

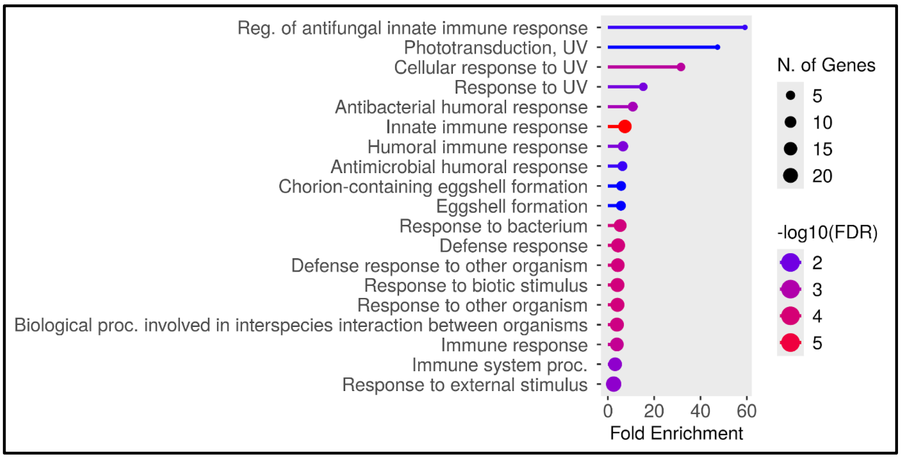


C

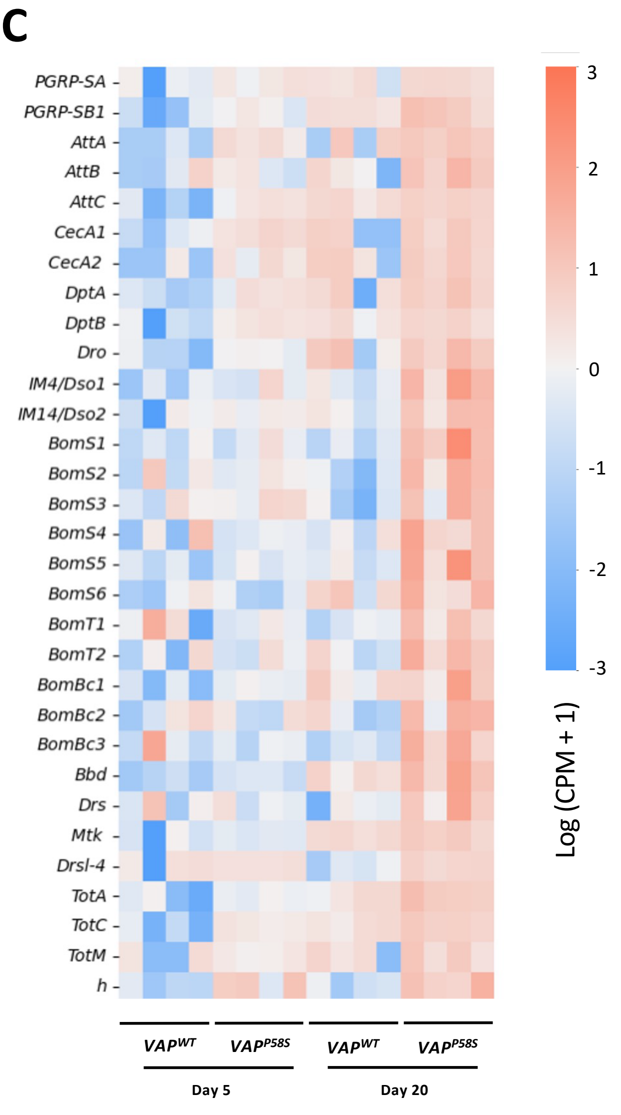


D

| | | | Ageing | | Disease | |
|---|---|---|---|---|---|---|
| | | | 20d vs 5d | | 5d | 20d |
| | Flybase ID | Flybase Symbol | *VAP^WT* | *VAP^P58S* | *VAP^P58S* vs *VAP^WT* | |
| Imd | FBgn0030310 | *PGRP-SA* | 0.81 | | 0.80 | 0.83 |
| | FBgn0043578 | *PGRP-SB1* | 1.89 | 1.93 | 1.19 | 1.79 |
| | FBgn0012042 | *AttA* | 1.03 | | | 1.75 |
| | FBgn0041581 | *AttB* | | 1.23 | | 1.39 |
| | FBgn0041579 | *AttC* | 3.47 | 1.59 | 2.48 | 1.91 |
| | FBgn0000276 | *CecA1* | 1.21 | 1.15 | 2.34 | 1.13 |
| | FBgn0000277 | *CecA2* | 1.31 | 1.60 | | 1.60 |
| | FBgn0004240 | *DptA* | 2.11 | 1.13 | 2.12 | 1.23 |
| | FBgn0034407 | *DptB* | 2.16 | 1.65 | 2.04 | 1.42 |
| | FBgn0010388 | *Dro* | 2.03 | 1.55 | 1.01 | 1.80 |
| Toll | FBgn0040653 | *IM4/Dso1* | | 0.80 | | 0.93 |
| | FBgn0067905 | *IM14/Dso2* | | 0.86 | | 0.97 |
| | FBgn0034329 | *BomS1* | | 1.01 | | 1.00 |
| | FBgn0025583 | *BomS2* | | 0.79 | | 0.83 |
| | FBgn0040736 | *BomS3* | | | | |
| | FBgn0034330 | *BomS4* | | 0.62 | | 0.64 |
| | FBgn0040734 | *BomS5* | | 1.21 | | 1.24 |
| | FBgn0040733 | *BomS6* | 0.70 | 0.95 | | 1.14 |
| | FBgn0262838 | *BomT1* | -0.95 | 1.17 | -0.81 | 1.16 |
| | FBgn0040735 | *BomT2* | | 1.07 | | 1.06 |
| | FBgn0034328 | *BomBc1* | 1.14 | 0.89 | | 1.35 |
| | FBgn0034331 | *BomBc2* | | 0.70 | | 0.68 |
| | FBgn0040582 | *BomBc3* | -0.77 | 1.38 | | 1.26 |
| | FBgn0034512 | *Bbd* | 0.98 | 1.21 | | 1.26 |
| | FBgn0283461 | *Drs* | -0.76 | 0.78 | | 1.03 |
| | FBgn0014865 | *Mtk* | 2.06 | 3.06 | | 2.86 |
| | FBgn0052282 | *Drsl-4* | -1.46 | 1.24 | | 1.30 |
| Jak/ STAT | FBgn0028396 | *TotA* | 1.85 | 2.46 | 0.72 | |
| | FBgn0044812 | *TotC* | 2.73 | | 2.08 | 2.05 |
| | FBgn0031701 | *TotM* | | 1.37 | | 1.27 |
| TF | FBgn0001168 | *h* | | | 1.20 | |

| -1.5 to -2 | -1 to -1.5 | -0.5 to -1 | 0.5 to 1 | 1 to 1.5 | 1.5 to 2 | 2-2.5 | 2.5-3 | 3-3.5 |
|---|---|---|---|---|---|---|---|---|

E

F

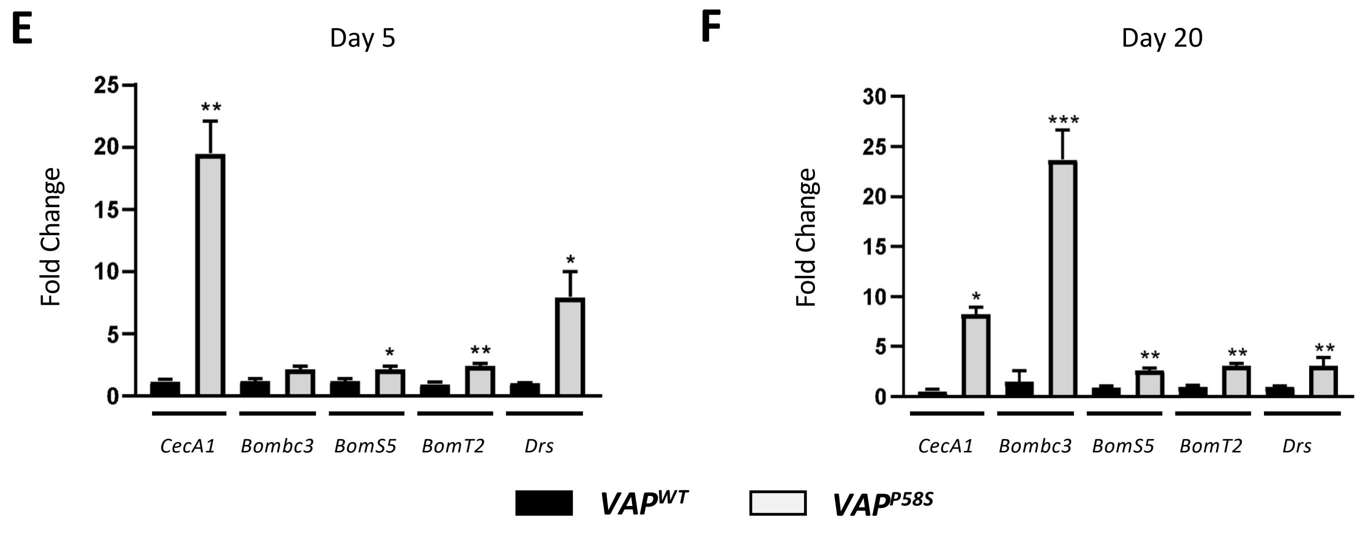

**Supplementary Figure 3: Transcriptomics on *VAP$^{P58S}$* female heads showing enhanced age-dependent neuroinflammation**

**A.** Genotypes and timepoints used for the 3' mRNA sequencing and the pairwise comparison done to obtain Differentially Expressed Genes (DEGs).

**B.** Gene Ontology (GO) enrichment analysis for significantly differentially expressed genes between *VAP$^{P58S}$* and *VAP$^{WT}$* at days 5 and 20 combined

**C.** Normalized expression counts for immune genes significantly differentially expressed between *VAP$^{P58S}$* and *VAP$^{WT}$* ,5d and 20d combined, depicted in the form of a heatmap

**D.** Summary table for immune gene transcripts categorized based on pathway/molecular function. Values represented as $log_2FC$ with the colour code given at the bottom.

**E & F.** qRT-PCR for the Imd pathway target *CecA1* and Toll pathway targets *Bombc3*, *BomS5, BomT2* and *Drs* at days 5 and 20 respectively for *VAP$^{WT}$* and *VAP$^{P58S}$* heads. The Y-axis shows $log_2FC$ values normalised to the housekeeping gene *rp49*. Values shown as mean + SEM. n= 40-50, N=3. Two-way ANOVA was used for statistical analysis. Tukey's test was used for multiple comparison. *$p< 0.05$, **$p< 0.01$, ***$p< 0.001$.

Suppl. Figure 4

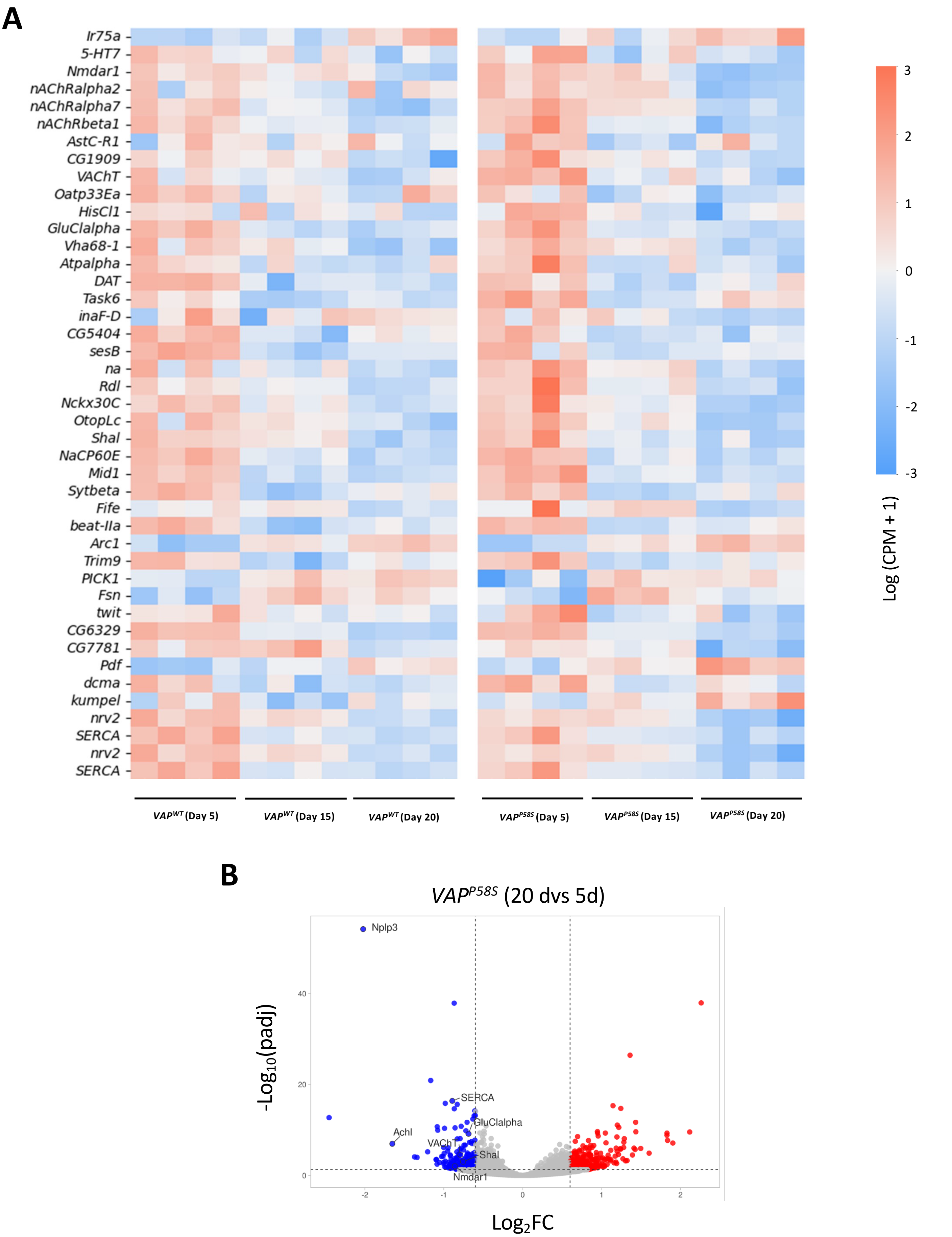


**Supplementary Figure 4: Heatmap and Volcano plot for *VAP*$^{P58S}$ male aging**

**A.** Heatmap showing the normalized expression counts for synaptic genes across genotypes and timepoints

**B.** Volcano plots representing up and downregulated genes for the complete dataset for *VAP*$^{P58S}$, 20d vs 5d. A few downregulated synaptic genes are highlighted

**A** *Up regulated*

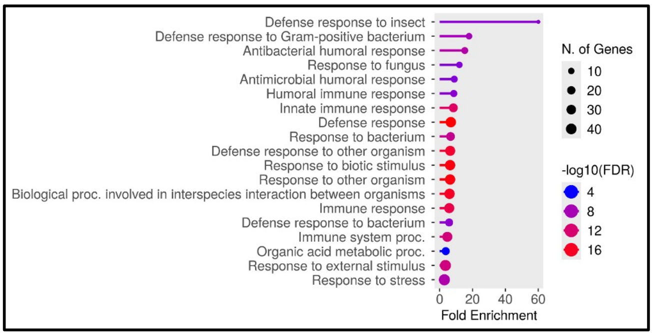


**B** *Down regulated*

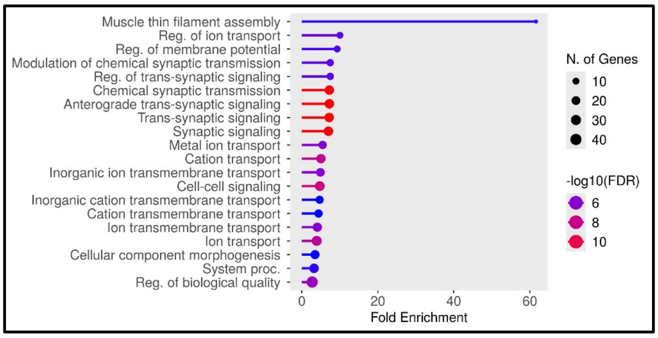


**C**

| Category | Flybase ID | Flybase Symbol | $VAP^{WT}$ 20d | $VAP^{P58S}$ 20d |
|---|---|---|---|---|
| Neurotransmitter receptors | FBgn0024944 | *Oamb* | 0.98 | |
| | FBgn0086778 | *nAChRalpha7* | | -1.01 |
| | FBgn0000038 | *nAChRbeta1* | | -0.95 |
| | FBgn0011582 | *Dop1R1* | 0.63 | |
| Transporters and Ion channels | FBgn0033196 | *HisT* | | -0.63 |
| | FBgn0024963 | *GluClalpha* | | -0.66 |
| | FBgn0003380 | *Sh* | | -0.69 |
| | FBgn0261794 | *kcc* | | -0.80 |
| | FBgn0259994 | *OtopLa* | | -0.64 |
| | FBgn0259145 | *CG42260* | | -0.66 |
| | FBgn0004244 | *Rdl* | | -0.65 |
| | FBgn0028704 | *Nckx30C* | | -0.64 |
| | FBgn0039915 | *Gat* | -0.63 | -0.69 |
| | FBgn0259150 | *OtopLc* | 0.68 | |
| | FBgn0039081 | *Irk2* | -0.70 | -0.82 |
| | FBgn0085434 | *NaCP60E* | | -0.82 |
| Synaptic vesicle release machinery | FBgn0004242 | *Syt1* | | -0.67 |
| | FBgn0033876 | *Syngr* | | -0.74 |
| | FBgn0032901 | *sky* | | -0.62 |
| Synapse formation and plasticity | FBgn0038498 | *beat-IIa* | | -0.96 |
| | FBgn0033926 | *Arc1* | | 0.74 |
| | FBgn0038975 | *Nrx-1* | | -0.65 |
| | FBgn0030174 | *CG15312* | 0.65 | -0.72 |
| | FBgn0040505 | *Alk* | | -0.63 |
| | FBgn0267001 | *Ten-a* | | -0.67 |
| Synaptic signalling and Modulators | FBgn0029663 | *CG10804* | | -0.71 |
| | FBgn0036007 | *path* | | 0.66 |
| | FBgn0263006 | *SERCA* | -1.11 | -0.77 |
| | FBgn0030897 | *Frq1* | -0.61 | -0.71 |

| | | | | | | | | |
|---|---|---|---|---|---|---|---|---|
| -1.5 to -2 | -1 to -1.5 | -0.5 to -1 | 0.5 to 1 | 1 to 1.5 | 1.5 to 2 | 2-2.5 | 2.5-3 | 3-3.5 |

**Supplementary Figure 5: Transcriptomic study in *VAP*$^{P58S}$ females showing changes in synaptic signalling with age**

**A.** Gene Ontology (GO) enrichment analysis for significantly upregulated genes in *VAP*$^{P58S}$ between days 5 and 20

**B.** Gene Ontology (GO) enrichment analysis for significantly downregulated genes in *VAP*$^{P58S}$ between days 5 and 20

**C.** Tabular representation of $log_2FC$ values for synaptic gene transcripts. The genes have been categorized into the functions they perform. The table shows comparison of *VAP*$^{P58S}$ at day 20 with *VAP*$^{P58S}$ at day 5 . The colour code is given at the bottom.

Suppl. Figure 5

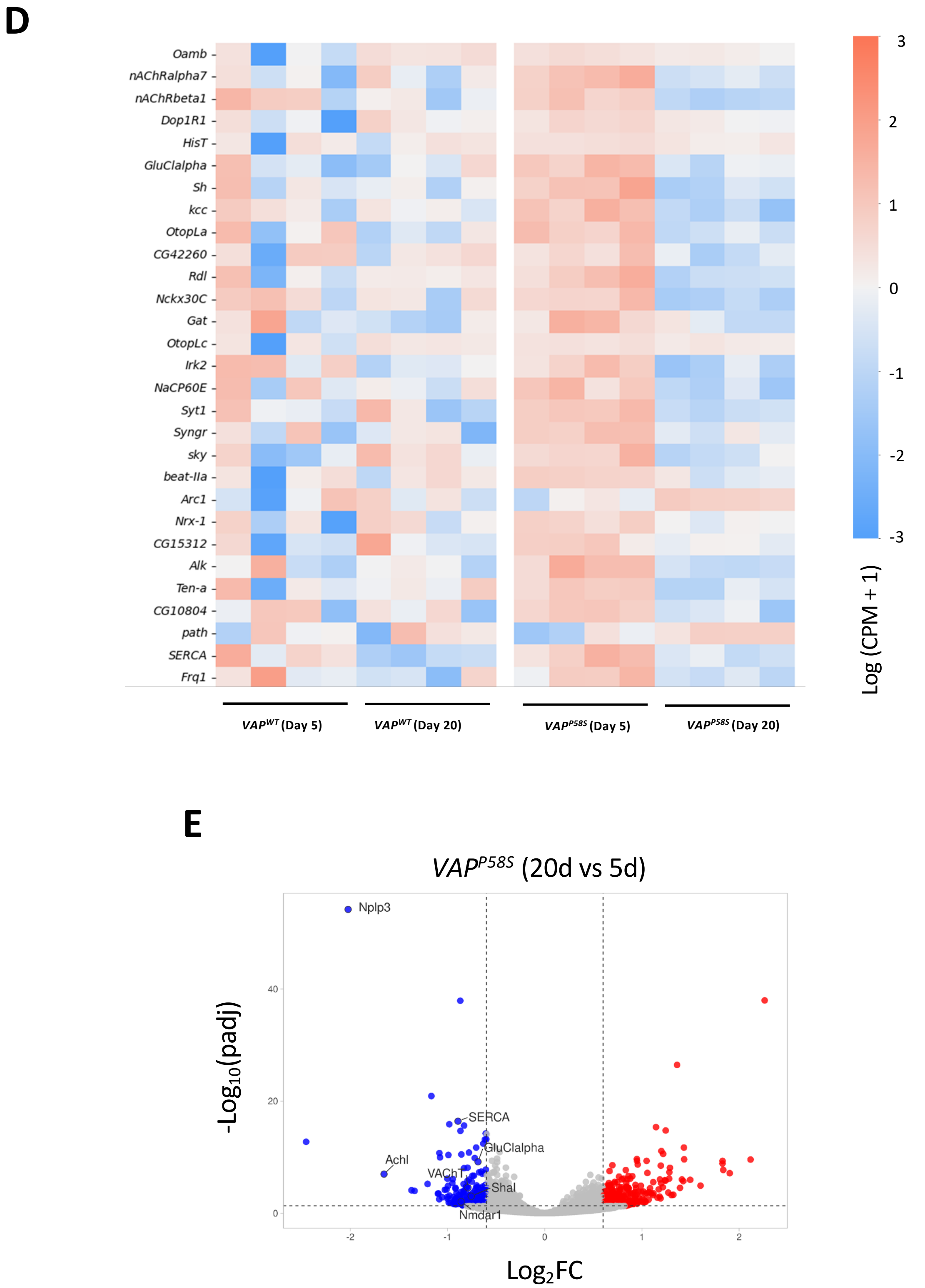


**D.** Heatmap showing the normalized expression counts for synaptic genes across genotypes and timepoints

**B.** Volcano plots representing up and downregulated genes for the complete dataset for *VAP*$^{P58S}$, 20d vs 5d. A few downregulated synaptic genes are highlighted

Suppl. Figure 6

**A**

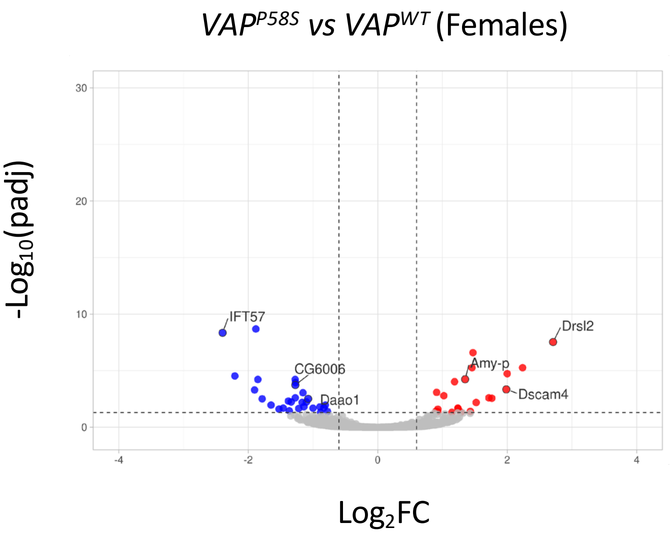


**B**

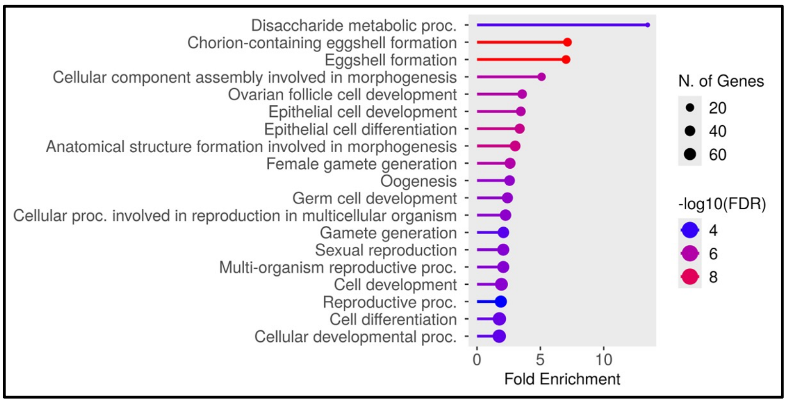


**C**

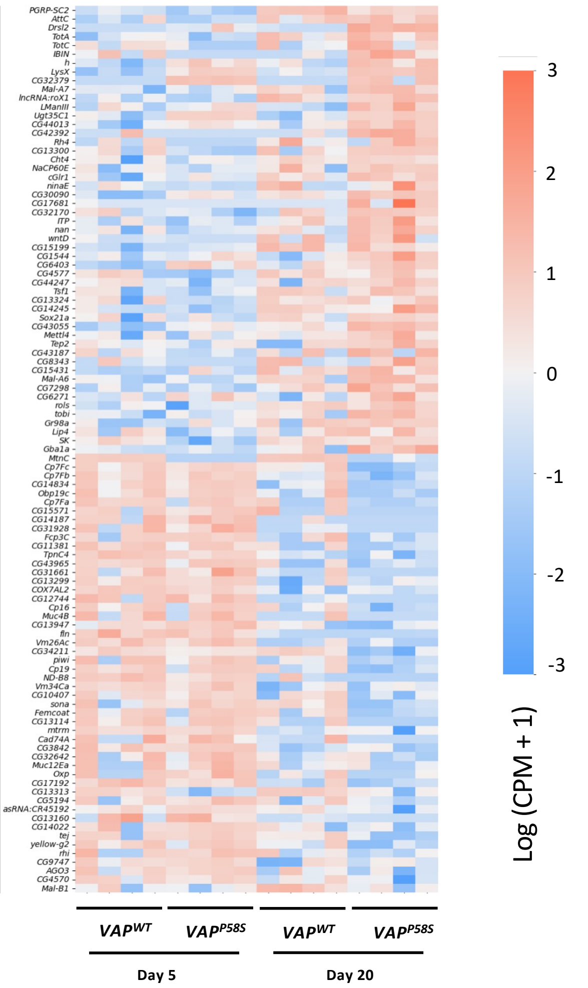


**D**

| | | Ageing | | Disease | |
|---|---|---|---|---|---|
| | | 20d vs 5d | | 5d | 20d |
| Flybase ID | Flybase Symbol | $VAP^{WT}$ | $VAP^{P58S}$ | $VAP^{P58S}$ vs $VAP^{WT}$ | |
| FBgn0043575 | *PGRP-SC2* | | | | 0.62 |
| FBgn0041579 | *AttC* | | 2.06 | | |
| FBgn0052279 | *Drsl2* | | 1.74 | | 2.71 |
| FBgn0028396 | *TotA* | | 1.92 | | |
| FBgn0044812 | *TotC* | | 1.47 | | |
| FBgn0265577 | *IBIN* | | 1.80 | | |
| FBgn0001168 | *h* | | | 1.29 | 1.04 |

| | | | | | | | | | | |
|---|---|---|---|---|---|---|---|---|---|---|
| -1.5 to -2 | -1 to -1.5 | -0.5 to -1 | 0 to -0.5 | 0 to 0.5 | 0.5 to 1 | 1 to 1.5 | 1.5 to 2 | 2-2.5 | 2.5-3 | 3-3.5 |

**Supplementary Figure 6: Gut transcriptomics for $VAP^{P58S}$ females**

**A.** Volcano plots for the genes of the entire dataset with a few important genes being highlighted

**B.** Gene Ontology (GO) enrichment analysis for significantly differentially expressed genes between $VAP^{P58S}$ and $VAP^{WT}$

**C.** Subset of genes significantly differentially expressed between $VAP^{P58S}$ and $VAP^{WT}$ for days 5 and 20 combined, plotted as a heatmap showing normalized expression counts

**D.** Summary table for immune gene transcripts. Values represent $log_2FC$ with the colour code given at the bottom.

### Suppl. Figure 7

A

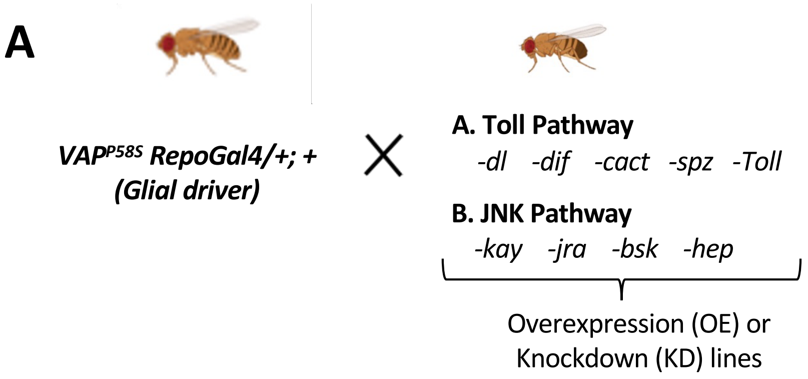


B

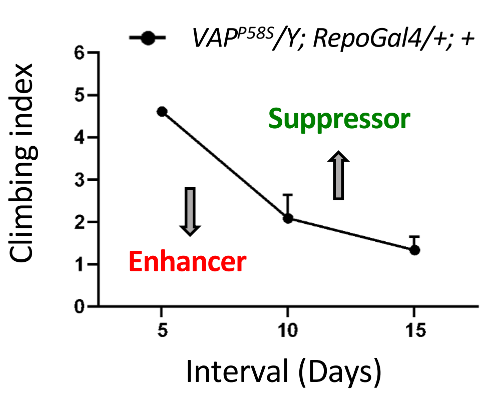


C

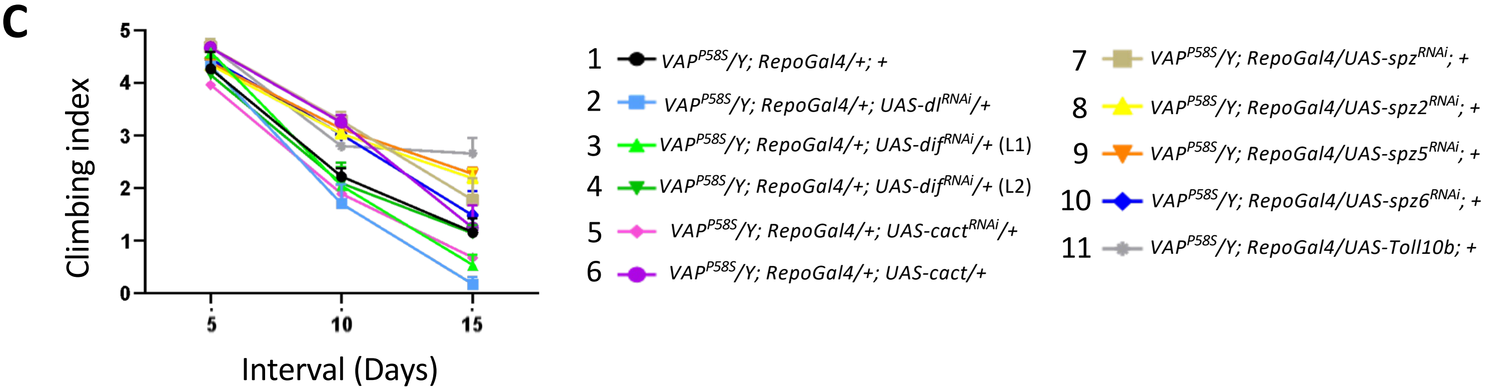


D

| | Toll pathway | P value for intervals | | |
|---|---|---|---|---|
| | Genotypes | 5 | 10 | 15 |
| 1 | *VAP^P58S/Y; RepoGal4/+; +* (Control) | | | |
| 2 | *VAP^P58S/Y; RepoGal4/+; UAS-dl^RNAi/+* | | | |
| 3 | *VAP^P58S/Y; RepoGal4/+; UAS-dif^RNAi/+* (L1) | | | |
| 4 | *VAP^P58S/Y; RepoGal4/+; UAS-dif^RNAi/+* (L2) | | | |
| 5 | *VAP^P58S/Y; RepoGal4/+; UAS-cact^RNAi/+* | | | |
| 6 | *VAP^P58S/Y; RepoGal4/+; UAS-cact/+* | | | |
| 7 | *VAP^P58S/Y; RepoGal4/UAS-spz^RNAi; +* | | | |
| 8 | *VAP^P58S/Y; RepoGal4/UAS-spz2^RNAi; +* | | | |
| 9 | *VAP^P58S/Y; RepoGal4/UAS-spz5^RNAi; +* | | | |
| 10 | *VAP^P58S/Y; RepoGal4/UAS-spz6^RNAi; +* | | | |
| 11 | *VAP^P58S/Y; RepoGal4/UAS-Toll10b; +* | | | |

Suppressor
- <0.0001 (****)
- <0.001 (***)
- <0.01 (**)
- <0.05 (*)
- >0.05

Enhancer
- <0.05 (*)
- <0.01 (**)
- <0.001 (***)
- <0.0001 (****)

E

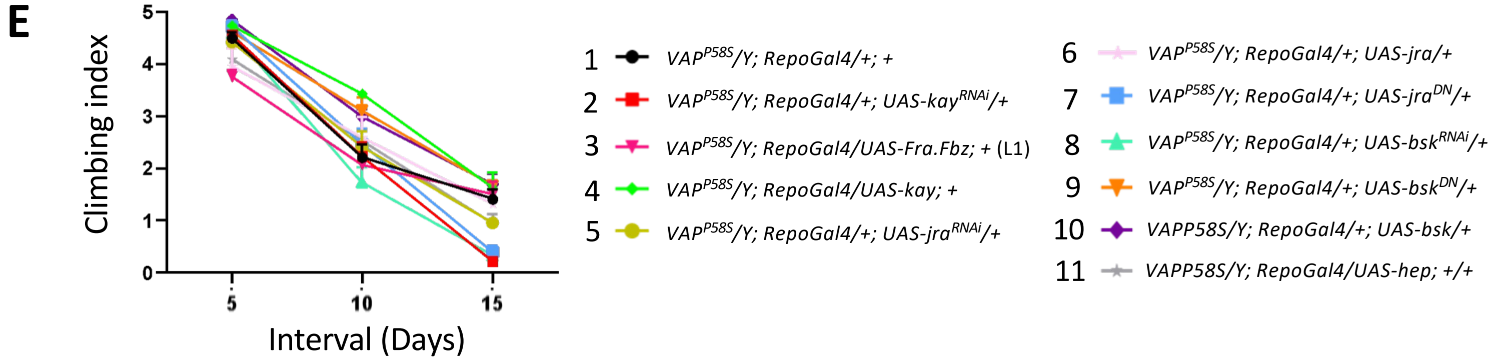


F

| | JNK pathway | P value for intervals | | |
|---|---|---|---|---|
| | Genotypes | 5 | 10 | 15 |
| 1 | *VAP^P58S/Y; RepoGal4/+; +* (Control) | | | |
| 2 | *VAP^P58S/Y; RepoGal4/+; UAS-kay^RNAi/+* | | | |
| 3 | *VAP^P58S/Y; RepoGal4/UAS-Fra.Fbz; +* (L1) | | | |
| 4 | *VAP^P58S/Y; RepoGal4/+; UAS-kay/+* | | | |
| 5 | *VAP^P58S/Y; RepoGal4/+; UAS-jra^RNAi/+* | | | |
| 6 | *VAP^P58S/Y; RepoGal4/+; UAS-jra^WT/+* | | | |
| 7 | *VAP^P58S/Y; RepoGal4/+; UAS-jra^DN/+* | | | |
| 8 | *VAP^P58S/Y; RepoGal4/+; UAS-bsk^RNAi/+* | | | |
| 9 | *VAP^P58S/Y; RepoGal4/+; UAS-bsk^DN/+* | | | |
| 10 | *VAP^P58S/Y; RepoGal4/+; UAS-bsk/+* | | | |
| 11 | *VAP^P58S/Y; RepoGal4/UAS-hep; +* | | | |
| 12 | *VAP^P58S/Y; RepoGal4/+; UAS-Fra.Fbz/+* (L2) | Lethal | | |
| 13 | *VAP^P58S/Y; RepoGal4/UAS-hep^CA; +* | Lethal | | |

**Supplementary Figure 7: Enhancer/Suppressor screen in the glia identified *kay* as an important regulator of motor activity in the *VAP$^{P58S}$* flies**

**A.** Experimental design showing glial knockdown or overexpression of various genes of Toll or JNK pathway using *VAP$^{P58S}$; RepoGal4; +*.

**B.** A schematic depicting the effect of gene modulation (knockdown or overexpression) on climbing index. The black curve represents the climbing index for *VAP$^{P58S}$/Y; RepoGal4/+; +* (control). A leftward shift indicates suppression (improved climbing), whereas a rightward shift indicates enhancement (worsened climbing) of the phenotype. The modulated gene is considered as a "suppressor" or "enhancer" depending on the direction of the shift.

**C.** Climbing indices for glial knockdown of *dl* (2, light blue), *dif* (L1) (3, light green), *dif* (L2) (4, dark green), *cact* (5, pink), *cact* overexpression (6, purple), knockdown of *spz* (7, brown), *spz2* (8, yellow), *spz5* (9, orange), *spz6* (10, dark blue) and *Toll10b* overexpression (11, grey). *VAP$^{P58S}$/Y; RepoGal4/+; +* (1, black) has been used as control. Values shown as mean $\pm$ SEM. n=25-30, N=3

**D.** Summary table for the gene modulations in panel (C) with the individual p value for each interval (day 5, 10, 15) depicted using a colour code. Statistical analysis was done by two-way ANOVA. Multiple comparison was done using Tukey's test. *$p<0.05$, **$p<0.01$, ***$p<0.001$, ****$p<0.0001$

**E.** Climbing indices for glial knockdown of *kay* (2, red), overexpression of *Fra.Fbz* (3, dark pink), overexpression of *kay* (4, light green), *jra* knockdown (5, ochre), *jra* overexpression (6, light pink), *jra$^{DN}$* overexpression (7, blue), knockdown of *bsk* (8, sea green), *bsk$^{DN}$* overexpression (9, orange), *bsk* overexpression (10, purple) and *hep* overexpression (11, grey). *VAP$^{P58S}$/Y; RepoGal4/+; +* (1, black) has been used as control. Values represented as mean $\pm$ SEM. n=25-30, N=3.

**F.** Summary table for the gene modulations in panel (E) with individual p values being listed. The colour scheme and statistical analysis is as previously described.

Suppl. Figure 8

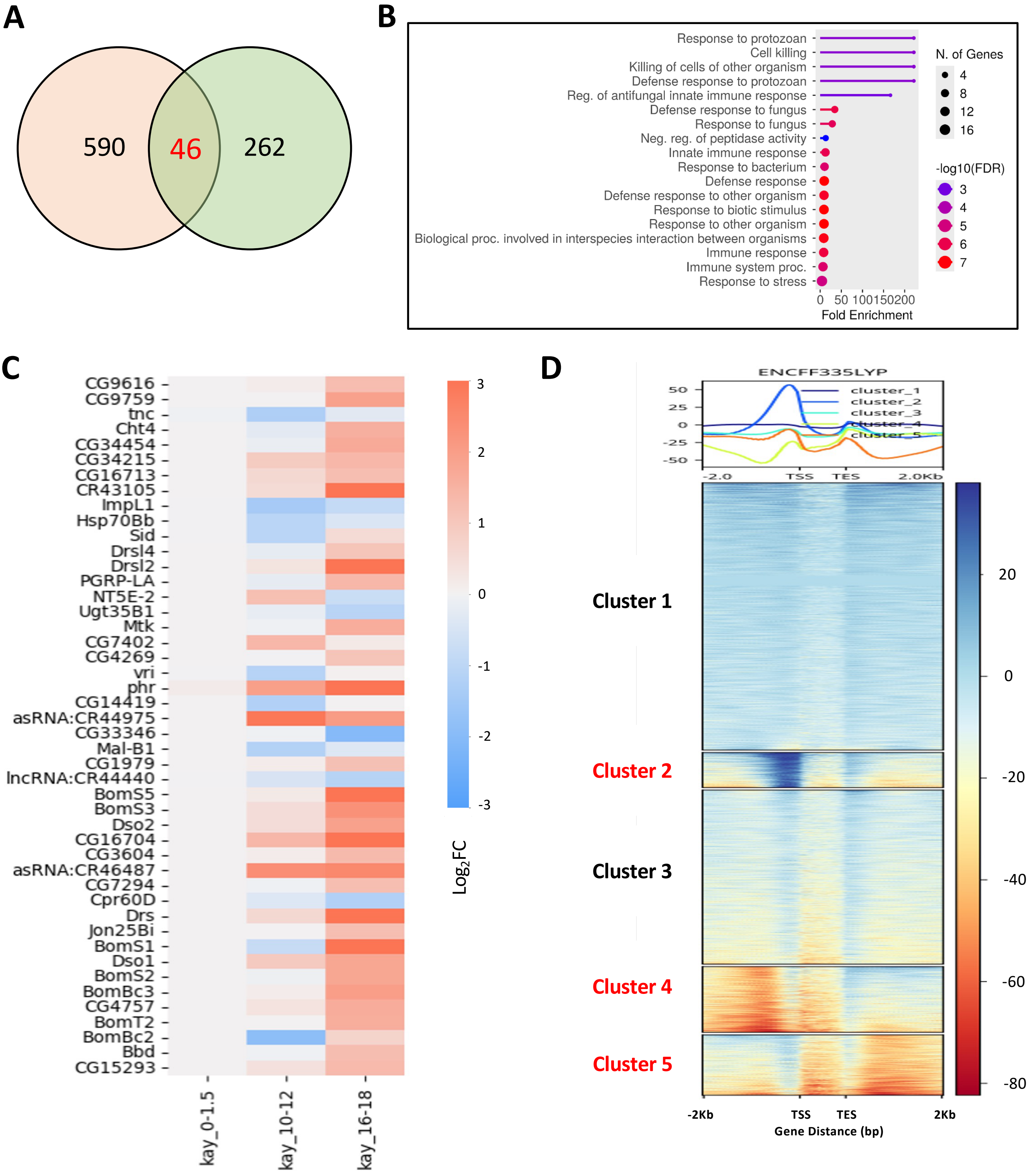


**E**

| Flybase ID | Flybase Symbol | Day | Fold Change (vs $VAP^{P58S}$; Repo/+; + (2)) | | |
|---|---|---|---|---|---|
| | | | 2 | 3 | 4 |
| FBgn0014018 | *Rel* | Day 5 | 1.42 | 0.96 | 0.96 |
| | | Day 15 | 1.30 | 0.74 | 0.86 |
| FBgn0260632 | *Dl* | Day 5 | 0.91 | 0.77 | 0.82 |
| | | Day 15 | 1.16 | 0.98 | 1.07 |
| FBgn0016917 | *STAT92E* | Day 5 | 0.84 | 0.97 | 1.01 |
| | | Day 15 | 0.87 | 0.95 | 0.82 |

*1: $VAP^{P58S}$/Y; RepoGal4/+; +*
*4: $VAP^{P58S}$/Y; RepoGal4/UAS-$Kay^{RNAi}$; +*
*3: $VAP^{P58S}$/Y; RepoGal4/UAS-Kay; +*
*4: $VAP^{P58S}$/Y; RepoGal4/UAS-$Kay^{SCR}$; +*

**Supplementary Figure 8: Kay regulates AMPs likely via an indirect mechanism**

**A.** Venn diagram showing overlap between the differentially expressed genes between *VAP*$^{P58S}$ and *VAP*$^{WT}$ (green circle) and the differentially expressed genes upon *kay* knockdown in the embryo (pink circle)

**B.** Gene Ontology (GO) analysis for the common genes in (A)

**C.** Heatmap showing the $log_2FC$ values for the common genes in (A) at various timepoints following *kay* knockdown in the embryo

**D.** Heatmap showing the occupancy of Kay on the gene body of all *Drosophila* genes. Clusters 2, 4 and 5 show genes with enriched binding of Kay. Data extracted from Kay-Chip Seq (ENCSR269HHK) against the input dataset (ENCSR331XEF)

**E.** Fold change for major immune pathway transcription factors upon glial *kay* modulation in the background of *VAP*$^{P58S}$

### Suppl. Figure 9

**A**

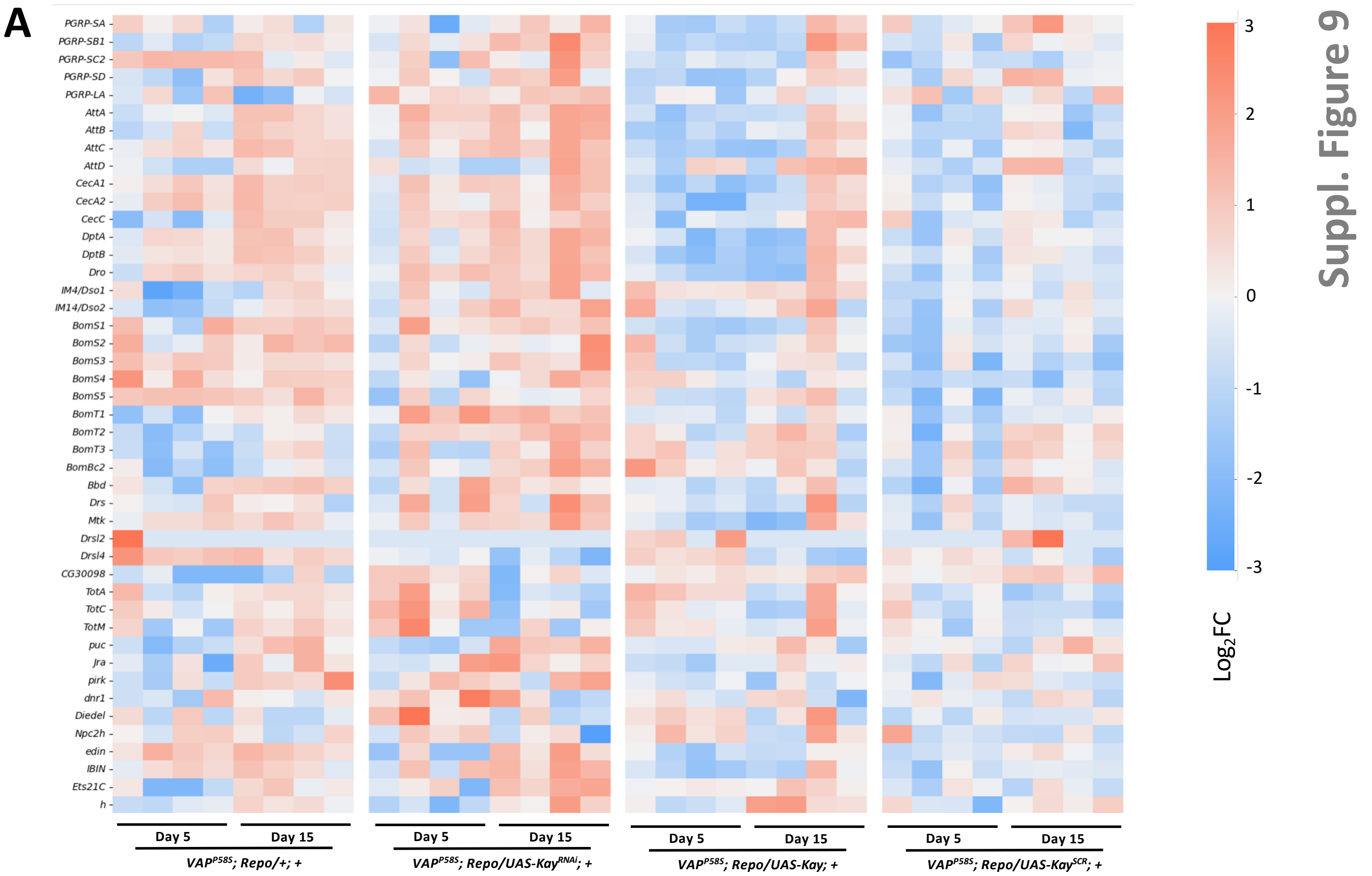


**B**

| Pathway | Flybase ID | Flybase Symbol | 1 | 2 | 3 | 4 |
|---|---|---|---|---|---|---|
| Imd | FBgn0030310 | *PGRP-SA* | | | | |
| | FBgn0043578 | *PGRP-SB1* | 0.74 | 0.86 | 1 | |
| | FBgn0043575 | *PGRP-SC2* | | | | |
| | FBgn0035806 | *PGRP-SD* | 0.61 | | 0.74 | |
| | FBgn0035975 | *PGRP-LA* | | | | |
| | FBgn0012042 | *AttA* | 1.14 | 0.62 | 0.94 | |
| | FBgn0041581 | *AttB* | 0.8 | 0.67 | 1.31 | |
| | FBgn0041579 | *AttC* | | 0.73 | | |
| | FBgn0038530 | *AttD* | | 0.69 | | 0.65 |
| | FBgn0000276 | *CecA1* | | | 0.6 | |
| | FBgn0000277 | *CecA2* | | | | |
| | FBgn0000279 | *CecC* | 0.72 | | 0.67 | |
| | FBgn0004240 | *DptA* | | 0.88 | | |
| | FBgn0034407 | *DptB* | | 0.7 | 0.8 | |
| | FBgn0034407 | *Dro* | | | 0.75 | |
| Toll | FBgn0040653 | *IM4/Dso1* | | | | |
| | FBgn0067905 | *IM14/Dso2* | | | | |
| | FBgn0034329 | *BomS1* | | | | |
| | FBgn0025583 | *BomS2* | | | | |
| | FBgn0040736 | *BomS3* | | | | |
| | FBgn0000276 | *BomS4* | | | | |
| | FBgn0040734 | *BomS5* | | | | |
| | FBgn0262838 | *BomT1* | 0.73 | | | |
| | FBgn0040735 | *BomT2* | | | | |
| | FBgn0038930 | *BomT3* | | | | |
| | FBgn0034331 | *BomBc2* | | | | |
| | FBgn0034512 | *Bbd* | | | | |
| | FBgn0283461 | *Drs* | | | 0.62 | |
| | FBgn0014865 | *Mtk* | | 0.6 | | |
| | FBgn0052279 | *Drsl2* | | | | |
| | FBgn0052282 | *Drsl4* | | -0.63 | -1.14 | |
| | FBgn0050098 | *CG30098* | | -0.62 | | 0.69 |
| Jak/STAT | FBgn0028396 | *TotA* | | -1.24 | | |
| | FBgn0044812 | *TotC* | | | | |
| | FBgn0031701 | *TotM* | | -1.2 | | |
| JNK | FBgn0243512 | *puc* | 0.64 | 0.76 | | |
| | FBgn0001291 | *Jra* | 0.74 | | | |
| IMs | FBgn0034647 | *pirk* | 0.78 | | | |
| | FBgn0260866 | *dnr1* | | | | |
| | FBgn0039666 | *Diedel* | | -1.19 | | |
| | FBgn0039801 | *Npc2h* | | | | |
| | FBgn0052185 | *edin* | | 0.92 | | |
| | FBgn0265577 | *IBIN* | | 0.85 | | |
| TFs | FBgn0005660 | *Ets21C* | 0.64 | 1.03 | | |
| | FBgn0001168 | *h* | | 0.7 | 0.62 | |

Colour key: 3-3.5; 2.5-3; 2-2.5; 1.5 to 2; 1 to 1.5; 0.5 to 1; -0.5 to -1; -1 to -1.5; -1.5 to -2

1: *VAP^P58S/Y; RepoGal4/+; +*
2: *VAP^P58S/Y; RepoGal4/UAS-Kay^RNAi; +*
3: *VAP^P58S/Y; RepoGal4/UAS-Kay; +*
4: *VAP^P58S/Y; RepoGal4/UAS-Kay^SCR; +*

**Supplementary Figure 9: Effects of *kay* modulation on age-dependent neuroinflammation in *VAP*$^{P58S}$**

**A.** Heatmap showing the normalized expression counts for the immune genes following *kay* modulation in the glia in the background of *VAP*$^{P58S}$

**B.** Tabular representation of $log_2FC$ values for immune gene transcripts with age following glial kay modulation in *VAP*$^{P58S}$ background

**Supplementary Table 1:** Details of the fly lines used in this study

| Sr. No. | Line | Genotype | Source | Details |
|---|---|---|---|---|
| 1 | *VAP$^{WT}$* | *VAP$^{WT}$; +; +* | In-house | VAP locus modified using CRISPR /Cas9. No mutation incorporated. Used as control |
| 2 | *VAP$^{P58S}$* | *VAP$^{P58S}$; +; +* | In-house | VAP locus modified using CRISPR/Cas9. VAP$^{P58S}$ mutation incorporated |
| 3 | *nos-Cas9* | v$^{-}$; *nanos-Cas9* | Gift from Ryu Ueda | |
| 4 | *VAP$^{P58S}$; RepoGal4/CyO; +* | *VAP$^{P58S}$; RepoGal4/CyO; +* | Balanced in-house | *VAP$^{P58S}$* balanced with glial Gal4 |
| 5 | *RepoGal4* | *+; RepoGal4/CyO; +* | Dr. Bradley Jones, University of Mississippi | Lee et al., 2005 |
| 6 | *UAS-dl$^{RNAi}$* | y[1] sc[*] v[1] sev[21]; P{y[+t7.7] v[+t1.8]=TRiP.HMS00727}attP2 | BDSC:32934 | |
| 7 | *UAS-dif$^{RNAi}$* (L1) | y[1] sc[*] v[1] sev[21]; P{y[+t7.7] v[+t1.8]=TRiP.HM05257}attP2 | BDSC:30513 | |
| 8 | *UAS-dif$^{RNAi}$* (L2) | y[1] v[1]; P{y[+t7.7] v[+t1.8]=TRiP.HM05191}attP2 | BDSC:29514 | |
| 9 | *UAS-cact$^{RNAi}$* | y[1] sc[*] v[1] sev[21]; P{y[+t7.7] v[+t1.8]=TRiP.HMS00084}attP2 | BDSC:34775 | |
| 10 | *UAS-cact* | *UASp-cact-eGFP* | Shubha Govind | |
| 11 | *UAS-spz$^{RNAi}$* | y[1] v[1]; P{y[+t7.7] v[+t1.8]=TRiP.HMJ22258}attP40 | BDSC:58499 | |
| 12 | *UAS-spz2$^{RNAi}$* | | VDRC:110495 | |
| 13 | *UAS-spz5$^{RNAi}$* | y[1] sc[*] v[1] sev[21]; P{y[+t7.7] v[+t1.8]=TRiP.HMC06330}attP40 | BDSC:67229 | |
| 14 | *UAS-spz6$^{RNAi}$* | y[1] sc[*] v[1] sev[21]; P{y[+t7.7] v[+t1.8]=TRiP.HMC04825}attP40 | BDSC:57510 | |
| 15 | *UAS-Toll10b* | P{w[+mC]=UAS-Tl.10b}11, y[1] w[*] | BDSC:58987 | Constitutively active Toll |

| | | | | |
|---|---|---|---|---|
| 16 | *UAS-kay$^{RNAi}$* | y[1] sc[*] v[1] sev[21]; P{y[+t7.7] v[+t1.8]=TRiP.HMS00254}attP2 | BDSC:33379 | *Drosophila* orthologue of Fos |
| 17 | *UAS-Fra.Fbz* (L1) | w[1118]; P{w[+mC]=UAS-Fra.Fbz}5 | BDSC:7214 | Dominant negative Kay |
| 18 | *UAS-Fra.Fbz* (L2) | y[1] w[1118]; P{w[+mC]=UAS-Fra.Fbz}7 | BDSC:7215 | Dominant negative Kay |
| 19 | *UAS-kay* (L1) | w[1118]; P{w[+mC]=UAS-Fra}2 | BDCS:7213 | |
| 20 | *UAS-jra$^{RNAi}$* | y[1] v[1]; P{y[+t7.7] v[+t1.8]=TRiP.JF01184}attP2 | BDSC:31595 | *Drosophila* orthologue of Jun |
| 21 | *UAS-jra* | y[1] w[1118]; P{w[+mC]=UAS-Jra}2 | BDSC:7216 | |
| 22 | *UAS-jra$^{DN}$* | y[1] w[1118]; P{w[+mC]=Jbz}10 | BDCS:7218 | Dominant negative Jra |
| 23 | *UAS-bsk$^{RNAi}$* | y[1] sc[*] v[1] sev[21]; P{y[+t7.7] v[+t1.8]=TRiP.HMS00777}attP2 | BDSC:32977 | |
| 24 | *UAS-bsk$^{DN}$* | w[*]; P{w[+mC]=UAS-bsk.K53R}20.1a | BDSC:9311 | Dominant negative Bsk |
| 25 | *UAS-bsk* | w[*]; P{w[+mC]=UAS-bsk.B}2 | BDSC:9310 | |
| 26 | *UAS-hep* | w[*]; P{w[+mC]=UAS-hep.B}2 | BDSC:9308 | |
| 27 | *UAS-hep$^{CA}$* | w[*]; P{w[+mC]=UAS-Hep.Act}2 | BDSC:9306 | Constitutively active Hep |
| 28 | *UAS-kay* (L2) | *+; UAS-kay$^{WT}$/CyO; +* | In-house | Inhouse |
| 29 | *UAS-kay$^{SCR}$* | *+; UAS-kay$^{SCR}$/CyO; +* | In-house | Inhouse-SUMO conjugation resistant form of Kay |
| 30 | *w[1118]* | | BDSC:3605 | |

**Supplementary Table 2:** Primers used in this study

| | | | |
|---|---|---|---|
| 1 | *pGEX-Kay F* | CTGGTTCCGCGTGGATCCCCGGAATTCATGA CGCTGGACAGCTACAAC | Primer pair used to amplify Kay for pGEX-4T1 |
| 2 | *pGEX-Kay R* | TCGTCAGTCAGTCACGATGCGGCCGCTTATA AGCTGACCAGCTTGGA | |
| 3 | *Kay-K357R F* | CTGATGCACATCAGGGACGAGCCACTC | Primer pair used to mutate K357R in Kay |
| 4 | *Kay-K357R R* | GAGTGGCTCGTCCCTGATGTGCAT | |
| 5 | *pRM-HA-Kay F* | ACGATGTTCCAGATTACGCTGGAGGCGAAAT GACGCTGGACAGCTACAA | Primer pair 28 pRM-HA-Kay R used to amplify Kay for pRM-HA3 |
| 6 | *pRM-HA-Kay R* | GCAGGTCGACTCTAGAGGATCCTTATAAGCT GACCAGCTTGG | |
| 7 | *pUASp-attB-Kay F* | GATCAGATCCGCGGCCGCATGTACCCATACG ATGTTC CAGATTACGCTGGAGGCGGAATGACGCTGG ACAGCTAC AAC | Primer pair to amplify Kay for pRM-HA3 |
| 8 | *pUASp-attB-Kay R* | TCTAGAGGATCCAGATCCACTAGTTTATAAG CTGACCAG CTTGGAC | |
| 9 | *pUASp-attB-HR-Kay F* | CGTTAGGTCCTGTTCATTGGTACCCGCCCGG GGATCAGA TCCGCGGCCGC | Primer pair to amplify pRM HA3 for Kay |
| 10 | *pUASp-attB-HR-Kay R* | CGTTAACGTTAACGTTCGAGGTCGACTCTAG AGGATCCA GATCCACTAGT | |
| 11 | *rp49 F* | *GACGCTTCAAGGGACAGTATC* | qRT-PCR primer to amplify *rp49* |
| 12 | *rp49 R* | *AAACGCGGTTCT GCATGAG* | |
| 13 | *CecA1 F* | *CAATCGGAAGCTGGGTG* | qRT-PCR primer to amplify *CecA1* |
| 14 | *CecA1 R* | *TAATCATCGG GTCAACCTCGGGC* | |
| 15 | *BomBc3 F* | *TCTGATCGGCGCTCATCCC* | qRT-PCR primer to amplify *BomBc3* |
| 16 | *BomBc3 R* | *TAGACGGGTTCACCATTCGG* | |
| 17 | *BomS5 F* | *TCAGTTTCATTTCAACCGTTGCC* | qRT-PCR primer to amplify *BomS5* |
| 18 | *BomS5 R* | *ATTTAACGGAGAAGCCACTGC* | |
| 19 | *BomT2 F* | *GGCAGCTGTTAATGCTACGC* | qRT-PCR primer to amplify *BomT2* |
| 20 | *BomT2 R* | *GGCTCCAGATGTGAGTGTGT* | |
| 21 | *Drs F* | *CTGTCCGGAAGATACAAGGG* | qRT-PCR primer to amplify *Drs* |
| 22 | *Drs R* | *TCGCACCAGCACT TCAGACT* | |
